# Variability in reported midpoints of (in)activation of cardiac I_Na_

**DOI:** 10.1101/2024.05.08.593173

**Authors:** Michael Clerx, Paul G.A. Volders, Gary R. Mirams

**Affiliations:** Centre for Mathematical Medicine & Biology, School of Mathematical Sciences, University of Nottingham, Nottingham NG7 2RD, UK; Department of Cardiology, Cardiovascular Research Institute Maastricht, Maastricht University Medical Center, Maastricht, 6202 AZ, The Netherlands

## Abstract

Electrically active cells like cardiomyocytes show variability in their size, shape, and electrical activity. But should we expect variability in the properties of their ionic currents? In this brief review we gather and visualise measurements of two important electrophysiological parameters: the midpoints of activation and inactivation of the cardiac fast sodium current, I_Na_. We find a considerable variation in reported mean values between experiments, with a smaller cell-to-cell variation within experiments. We show how the between-experiment variability can be decomposed into a correlated and an uncorrelated component, and that the correlated component is much larger and affects both midpoints almost equally. We then review biological and methodological issues that might explain the observed variability, and attempt to classify each as within-experiment or correlated and uncorrelated between-experiment factors. Although the existence of some variability in measurements of ionic currents is well-known, we believe that this is the first work to systematically review it and that the scale of the observed variability is much larger than commonly appreciated, which has implications for modelling and experimental design.

## Introduction

Variability in electrophysiological properties arises at several scales. Between and within subjects, electrically active cells such as cardiomyocytes and neurons vary in number (Olivetti et al., 1995), size and shape (Volders et al., 1998), and ion channel expression levels (Schulz et al., 2006). But as we continue down the scales, towards molecules and atoms and into the realms of chemistry and physics, we may expect *biological* variability to disappear.

Where do ion channels fit in this picture? Transcription, translation, anchoring, and degradation of ion channel genes can affect the total number of channels in a cell, and hence the maximal conductance of its aggregate (whole-cell) currents. But should we also expect cell-to-cell or intersubject differences in properties that are not governed by channel count, such as voltage-dependence? Ion channel function is known (or suspected) to be modulated by several mechanisms, including localisation, phosphorylation, stretch, and even proximity to other channels (Marionneau and Abriel, 2015; Daimi et al., 2022; Beyder et al., 2010; Hichri et al., 2020). But what is the impact of such mechanisms on variability in ‘baseline’ currents, measured under controlled experimental conditions?

Here, we address this question using literature data gathered for a previous study on the human cardiac fast sodium current, I_Na_(Clerx et al., 2018). Where our earlier study focused on mutants, here shall use exclusively the accompanying wild-type controls. To gain a large but uniform data set, we will focus on the most common experiment type in this database: measurements using the whole-cell patch-clamp configuration in cells heterologously expressing *SCN5A*, the primary subunit of the channels conducting I_Na_ in the heart. Similarly, although I_Na_ voltage-dependence is complex, we shall focus on two of the most common quantities used to characterise it: the midpoints of activation (*V*_*a*_) and inactivation (*V*_*i*_). These describe the voltage at which the channel is half-maximally activated (or the voltage at which the measured peak conductance is half the maximum observed value), and the voltage at which it is half-maximally inactivated (see e.g. Sakakibara et al., 1992; Chadda et al., 2017).

Each *experiment* in our data set consists of measurements of *V*_*a*_ and/or *V*_*i*_ in several cells, expressed as a mean (*µ*_*a*_ and *µ*_*i*_, respectively), a standard deviation (*σ*_*a*_ and *σ*_*i*_), and a cell count (*n*_*a*_ and *n*_*i*_). We shall use the individual standard deviations as a measure of *within-experiment* variability, and refer to the difference between the means as *between-experiment* variability.

## Methods

All data used in this study were collected as part of a previous study on single-point mutations in *SCN5A* in expression systems (Clerx et al., 2018). For the current study, we reduced this data set to keep only wild-type (control) measurements, and we removed Xenopus oocyte measurements to keep only whole-cell patch-clamp studies. The systematic process whereby the original data was gathered is detailed below. Although this is not a study into effect sizes, we followed the PRISMA guidelines (Page et al., 2021) where applicable.

To identify candidate studies, we searched PubMed for “SCN5A mutation” (with the last search occurring in May 2016) and looked in previously published lists of mutations (Napolitano et al., 2003; Moric et al., 2003; Ackerman et al., 2004; Zimmer and Surber, 2008; Hedley et al., 2009; Kapplinger et al., 2010, 2015). Studies identified this way were then scanned to see if they contained measurements of *V*_*a*_ or *V*_*i*_ made with whole-cell patch clamp in either HEK or CHO cells, along with the number of cells measured and a standard deviation or standard error of the mean. Next, we filtered out studies made at normal or raised body temperatures, but kept studies made at “room temperature” (as stated by the authors) or at any temperature in the range from 18 to 26^*°*^C. Because of the considerable effort involved in performing experiments at body temperature, we assumed that studies not mentioning temperature satisfied our criteria and could be included. Similarly, we excluded any studies under non-baseline conditions (e.g. with known stretch, remodelling, ischemia, etc.). All data collection and selection was performed by M.C.

The dataset includes measurements in two different expression systems: HEK293 or tsA201 (both indicated as “HEK” in this study), and CHO cells. A clear statement of cell type was part of the inclusion criteria (see above), so that no missing-data strategy was required. The exact *SCN5A α*-subunit expressed in these cells was not always clearly indicated. We found at least four different isoforms, which we labelled: **a**, sometimes known as Q1077 and with GenBank accession number AC137587; **b**, known as Q1077del or GenBank AY148488; **a\***, hH1, R1027Q, or GenBank M77235; and **b\***, hH1a or T559A; Q1077del, no GenBank number (see also Makielski et al., 2003). Missing *α*-subunit information was recorded as “*α*-subunit unknown”. Finally, we noted whether or not studies stated a co-expressed *β*1 subunit; no information on *β*1 co-expression was taken to mean it was not co-expressed.

Some studies we surveyed recorded separate control (wild-type) experiments for each mutant (see Supplementary Table 1). We therefore distinguish between *studies* and *experiments*, where a study can contain several experiments, and each experiment summarises findings in multiple cells.

For each experiment, we recorded either the midpoint of activation (as a mean *µ*_*a*_, a sample standard deviation *σ*_*a*_, and a cell count *n*_*a*_), the midpoint of inactivation (*µ*_*i*_, *σ*_*i*_, and *n*_*i*_), or both. Sample standard deviations were not usually given directly, but could be calculated from the provided standard errors of the mean (SEMs). In Figure 1 we shall make the further assumption that midpoints in individual cells were distributed normally, allowing us to plot a 5^th^-to-95^th^ percentile range of the corresponding normal distribution.

**Figure 1:**
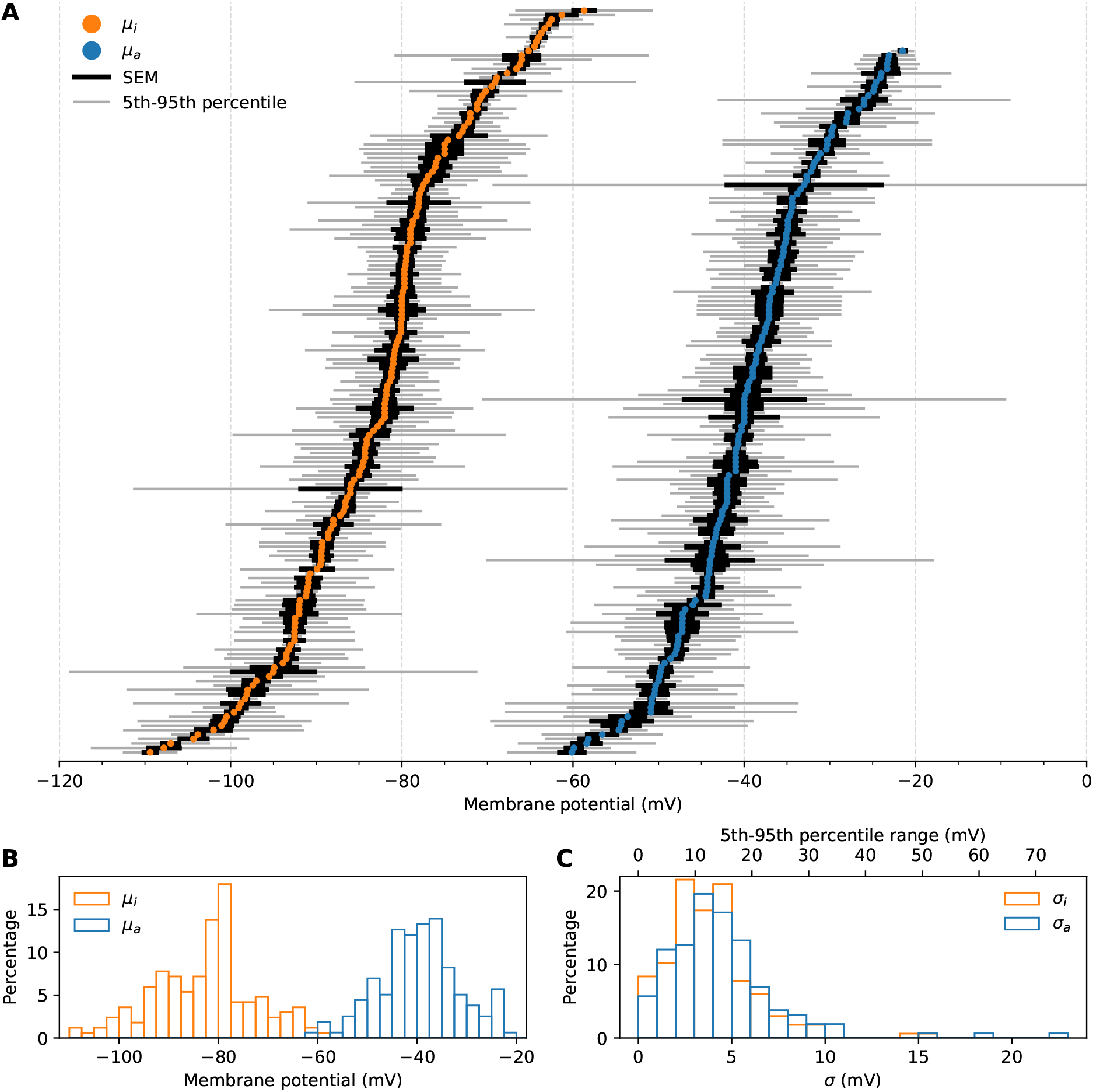
*A*, Reported mean midpoints of inactivation (*µ*_*i*_, left) and activation (*µ*_*a*_, right) for all experiments. Vertically, both sets of points are individually ordered from most to least negative membrane potential: correlations between an experiment’s *µ*_*a*_ and *µ*_*i*_ cannot be seen here and will be examined below. The standard error of the mean (SEM) for each experiment is indicated by a thick black bar. A thinner grey bar shows the 5^th^-to-95^th^ percentile range of a normal distribution with the reported mean and standard deviation: if the individual cell measurements in these studies were normally distributed, 90% of measurements would fall within this range. *B*, A histogram view of the means. The *y*-axis shows the percentage of reported means with each potential. *C*, A histogram view of the standard deviations. A second *x*-axis (top) shows the corresponding 5^th^-to-95^th^ percentile ranges.

Where both midpoints were reported, cell counts were often equal (34%) or similar (differing by no more than 5 cells in 90% of experiments, see Supplementary Table 2). So whilst it is plausible that *V*_*a*_ and *V*_*i*_ were often both measured in the same cell this cannot be guaranteed (and was not explicitly stated in many papers). However, we will assume that, even when cell counts were different, the conditions under which *V*_*a*_ and *V*_*i*_ were measured in an experiment were similar enough that correlations between *µ*_*a*_ and *µ*_*i*_ can be studied.

We shall use the term *within-experiment variability* to refer to the standard deviations within individual experiments, and the term *between-experiment variability* to refer to differences in reported means between different experiments (even in cases where those means are reported in the same study). For the two studies containing more than 5 experiments, we shall also look at *within-study between-experiment variability*, again by comparing the reported means.

## Data availability

The full list of midpoints and references can be found in Supplementary Table 3. A database version of the same data, along with code to generate all figures, tables, and numbers in this study, is available for download from https://github.com/MichaelClerx/ina-midpoints.

## Results

In the 120 studies that met the selection criteria, we found a total of 174 experiments: 151 experiments reporting both midpoints, 7 reporting only on activation, and 16 reporting only on inactivation. The obtained means (*µ*_*a*_ and *µ*_*i*_) and standard errors of the mean (SEM) are shown graphically in Figure 1. To see where the individual cell estimates of *V*_*a*_ and *V*_*i*_ may have been, for each experiment we also plot the 5^th^-to-95^th^ percentile range of a normal distribution with the reported *µ* and *σ* (approximately the range *µ ±* 1.64*σ*).

Within-experiment variability can be seen in the grey bars in panel A and the histograms in panel C. The median standard deviations were 3.6 mV for *V*_*i*_ and 4.0 mV for *V*_*a*_. Assuming a Normal distribution, this suggests that 90% of single-cell results in a typical experiment fall in a range of approximately 12 mV (*V*_*i*_) to 13 mV (*V*_*a*_). Slightly larger ranges of up to 20 mV or 30 mV are also not uncommon (panel C, top axis), but outliers go up to 50 mV (*V*_*i*_) and 73 mV (*V*_*a*_).

More surprisingly, substantial between-experimental variability can be seen in our data set: reported means of *V*_*i*_ range from *−*109 to *−*59 mV (median *−*81.4 mV, range 50.7 mV, 5^th^-to-95^th^ percentile range 35.5 mV), while means of *V*_*a*_ range from *−*60 to *−*21 mV (median *−*40.0 mV, range 38.6 mV, 5^th^-to-95^th^ percentile range 28.9 mV). Despite the large between-experiment variability, the SEM for most experiments, which quantifies the degree of certainty in the estimate of the mean, is quite narrow. This suggests that the mean *V*_*a*_ and *V*_*i*_ differed significantly between the surveyed experiments, and that one or more confounding factors may exist that explain this difference. Inactivation results seem more affected, which a much larger between-experiment variability for *µ*_*i*_, while the median within-experiment variability *σ*_*i*_ is slightly smaller than *σ*_*a*_.

### Mean midpoints *µ*_*a*_ and *µ*_*i*_ strongly correlate across experiments

Next, we look at *V*_*a*_ and *V*_*i*_ in the subgroup of 151 experiments where both were reported, as shown in Figure 2A. Each experiment is shown as a dot and a linear fit through all experimental means is shown, made using unweighted least-squares based linear regression. This line had an offset of *−*45.4 mV and a slope of 0.94 mV*/*mV, with a Pearson correlation coefficient *R* = 0.79. The coefficient of determination was *R*^2^ = 0.63, indicating that 63% of the variance is explained by this linear correlation. A second regression with a fixed slope of 1 is shown (green line), and this falls within the 95% confidence interval of the original regression (shaded grey area and dashed blue lines), so that we cannot statistically reject the hypothesis that the slope equals 1. Together, this correlation suggests the existence of some unknown factor shifting *µ*_*a*_ and *µ*_*i*_ by approximately equal amounts between experiments.

**Figure 2:**
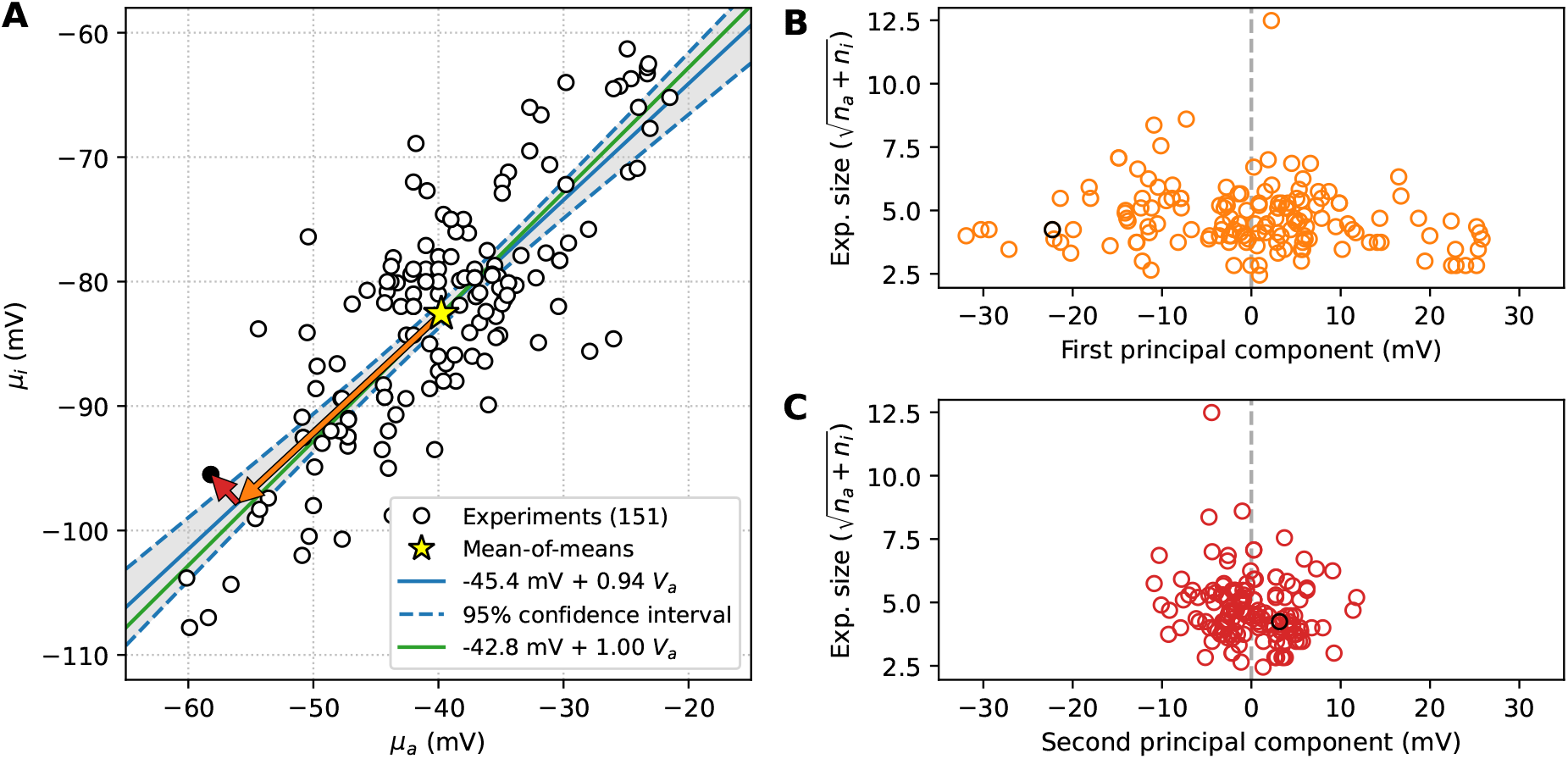
*A*, Mean midpoints of inactivation *µ*_*i*_ plotted against mean midpoints of activation *µ*_*a*_, for the experiments that reported both. The mean of all points (a mean-of-means) is indicated by a yellow star. A best-fit line is shown as a solid blue line, with its 95% confidence interval indicated by dashed blue lines and a gray shaded area. A second linear regression line with a slope constrained to have a gradient of one is shown in green. For one example experiment (*µ*_*a*_ = *−*58.2 mV, *µ*_*i*_ = *−*95.5 mV) we show the vector from the mean-of-means to this point, decomposed into components along the line of best fit (first principal component) and perpendicular to the line of best fit (second principal component). The same example point is highlighted in black in panels B and C. *B*, The square root of the experiment size as a function of the first principal component, for all points in A. The experiment size is defined as *n*_*a*_ + *n*_*i*_, where *n*_*a*_ is the number of cells tested for *V*_*a*_ and *n*_*i*_ is the number tested for *V*_*i*_. *C*, The square root of the experiment size as a function of the second principal component.

We can decompose the difference between each (*µ*_*a*_, *µ*_*i*_) measurement and the group mean into a component along the line of best fit (without constraining the slope), and a component perpendicular to the line of best fit (i.e. principal component analysis, PCA). An example for a single point is shown by the arrows drawn in Figure 2A), and the same example point is highlighted in black in panels B and C. The result suggests that most of the between-experiment variability is positively correlated.

In panels B and C we test whether the variability in either direction diminishes with experiment size (number of cells tested). To this end, we define ‘experiment size’ as the number of cells *n*_*i*_ tested to measure *µ*_*i*_, plus the number of cells *n*_*a*_ tested to measure *µ*_*a*_. In Figure 2B we plot the square root of this quantity 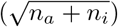 as a function of the first principal component to create something akin to a ‘funnel plot’. No clear triangle shape is observed in either plot, but the first component does appear to somewhat diminish with an increased number of measurements.

### Subunits and cell type are not the major sources of variability

Cell type, *α*-subunit isoform, and *β*1-subunit co-expression may affect *V*_*a*_ and *V*_*i*_ and were duly reported in most publications we checked. But can they explain the large between-experiment variability we observed? In Figure 3, we show the same data as in Figure 2, but grouped by recorded *α* subunit, *β*1 co-expression, and cell type. The largest subgroup (a* subunit, with *β*1 co-expression, in HEK) is shown in panel D. It is clear that, while some differences between these groups exist which could cause subtle shifts in the means, grouping like this does not divide our data into clear-cut clusters. In fact, many of the larger groups span the full observed range, suggesting that these factors have only a small effect on *V*_*a*_ and *V*_*i*_ measurements — even though their effect on *in vivo* electrophysiology may be profound.

**Figure 3:**
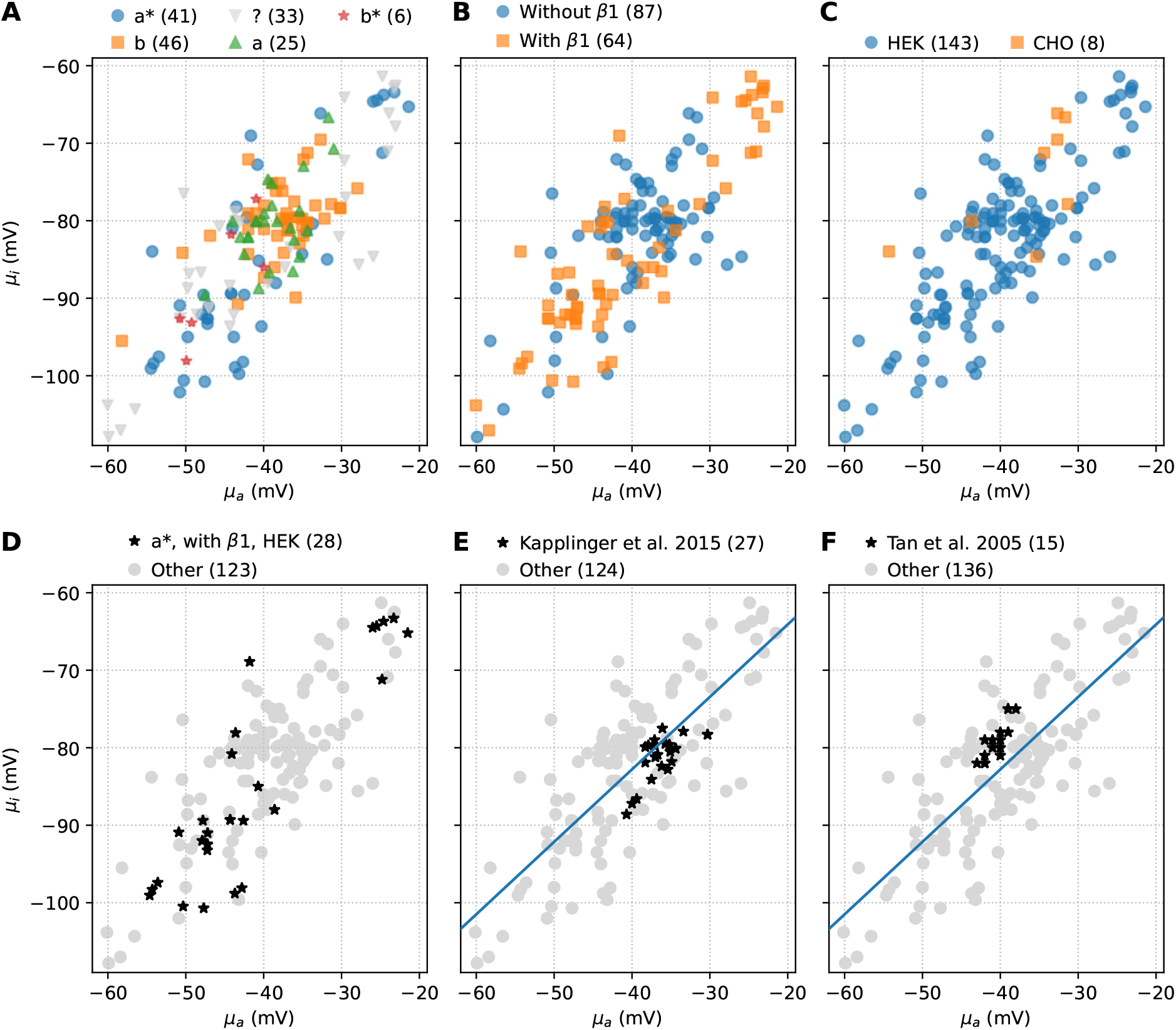
Grouping by recorded *α* subunit, *β*1 subunit co-expression, and cell type does not create distinct clusters and only explains a small part of the observed between-experiment variability. The number after each category indicates the corresponding number of means. Within-study between-experiment variability is observed in the two largest studies, but is much smaller than in the full data set. *A*, Grouping by *α* subunit; from largest to smallest subgroup we show the a* (R1027Q) *α* subunit, b (Q1077del), not reported, a (Q1077), and b* (T559A; Q1077del). *B*, Grouping by *β*1 co-expression. *C*, Grouping by cell type (HEK versus CHO cells), but note the very different group sizes. *D*, The largest subgroup versus all other results. *E*, Within-study variability in the study by Kapplinger et al. (2015) and *F*, Tan et al. (2005).

### Within-study between-experiment variability

The final two panels in Figure 3 show the two studies with more than five experiments: Kapplinger et al. (2015, 27 experiments), and or Tan et al. (2005, 15 experiments). Again, a strong correlated component is visible in both. Compared to the full data set, both correlated and uncorrelated between-experiment variability is much smaller in these groups.

## Discussion

We observed strong variability within experiments (median *σ*_*i*_ was 3.6 mV, median *σ*_*a*_ was 4.0 mV, but with outliers up to 22 mV) and between experiments (*µ*_*i*_ ranged from *−*109 to *−*59 mV, *µ*_*a*_ from *−*60 to *−*21 mV), and found a strong positive correlation across experiments measuring both (explaining 63% of the observed between-experiment variance). Cell type, *α*-subunit, and *β*1-subunit were seen to have an influence, but grouping by these categories did not explain the results. We also saw within-study between-experiment variability, on a smaller scale but with a visually similar correlation. How should we interpret these findings?

The existence of *some* within-experiment variability is well known, and is the reason why midpoints are reported as a mean and SEM. The existence of between-study or between-lab variability too, is indirectly acknowledged by the mutant studies we collected in Clerx et al. (2018) and reused here: each provided a new wild-type recording instead of using a value from the literature. Some studies measuring multiple mutants have gone even further, and accounted for *within-study between-experiment* variability by performing a paired control wild-type measurement for every measured mutant. For good examples see Kapplinger et al. (2015, 27 reported wild-type values) or Tan et al. (2005, 15 reported wild-type values). The Tan et al. (2005) paper also provides the only *direct* acknowledgement of between-experiment variability we found, citing “seasonal variation in current characteristics” as a reason for their paired study design. However, the wild-type values reported in Tan et al. (2005) and Kapplinger et al. (2015) differ by at most 11 mV and 7 mV respectively — well short of the 40 and 50 mV ranges seen in Figure 1. The full extent of between-experiment variability then, is still surprising.

Interestingly, the least negative (most depolarised) reported value of *µ*_*i*_ is -58.7 mV, exceeding the most negative (least depolarised) *µ*_*a*_ of -60.1 mV. Such a situation is clearly not physiological, and it is tempting to postulate some unknown biological mechanism (present even in cells non-natively expressing *SCN5A*) that regulates the difference between the midpoints, keeping *V*_*a*_ *− V*_*i*_ at approximately 45 mV and explaining the correlation with a gradient indistinguishable from 1 that is seen in Figure 2. However, a simpler explanation might be sought in experimental factors causing a difference between the intended and the applied voltage that applies equally to measurements of *V*_*a*_ and *V*_*i*_. We briefly review such factors below.

### Experimental sources of variability

An overview of experimental sources of variability (or more precisely, *uncertainty* that might cause variability in measurements, see Mirams et al., 2016) is shown in Table 1, and we have made a tentative effort to classify each as causing between- or within-experiment variability. The between-experiment column is further divided into correlated and uncorrelated effects. Disputed or hypothetical factors are indicated using question marks, while check marks indicate factors known to strongly influence results — although the extent of their effect on our data is still unknown. In the text below, we explain our reasoning and provide some highly speculative upper bounds on effect magnitudes.

**Table 1:** Postulated experimental causes of variability. Characterised as strongly likely (✓✓), likely (✓), possibly (?), or unlikely (✗) to be contributing to the different types of variability in measurements of *µ*_*a*_ and *µ*_*i*_.

|  | Between experiment |  | Within experiment |
| --- | --- | --- | --- |
|  | correlated | uncorrelated |  |
| Temperature | ? | ✗ | ? |
| Time since rupture | ? | ? | ? |
| Missing or erroneous LJP correction | ✓✓ | ✗ | ✗ |
| Voltage-control errors | ✓ | ✓ | ✓ |
| Voltage protocol and analysis | ✓ | ✓ | ✗ |
| Stretch | ✗ | ✗ | ? |
| Endogenous currents | ? | ? | ? |
| Regulation | ? | ? | ? |

The measurements we reviewed were made at “room temperature”, defined by the various authors as anywhere between 18^*°*^C and 26^*°*^C. Nagatomo et al. (1998) recorded a shift in the midpoint of activation of +0.43 mV per ^*°*^C, and a +0.47 mV shift for inactivation, although no such shifts were observed by Keller et al. (2005) and both studies used HEK cells. If there is a 0.5 mV per ^*°*^C shift, the observed range of room temperatures could lead to a correlated between-experiment effect of up to 4 mV. Within *studies* temperature was usually given as a 1 or 2 degree bracket, leading to a much smaller within-experiment estimate of 0.5 to 1 mV.

Hanck and Sheets (1992) measured I_Na_ in Purkinje cells and studied the effect of the time between rupturing the membrane and performing the measurement, which caused both midpoints to drift towards more negative potentials at approximately 0.5 mV per minute. A study by Abriel et al. (2001) looked for, but did not find evidence of, a similar time-dependent drift in HEK cells. Time since rupture was not reported in the studies we reviewed, which makes it difficult to classify this effect. First, between-experiment variability may arise if highly systematic approaches are employed but these differ between experiments/studies. Any unsystematic deviation cell-to-cell, e.g. due to the time needed to note down cell measurements or adjust compensation circuitry, will lead to within-experiment variability. Next, a correlated means effect could arise for example if a systematic approach was followed, if both midpoints were measured in the same cells (consistent with the similar *n*_*a*_ and *n*_*i*_ shown in Supplementary Table 1), and if the time between activation and inactivation protocols was short relative to the time needed to set up. Because of these uncertainties, we list “time since rupture” as only a possible effect in all three columns of Table 1. The magnitude of these three effects is impossible to determine from our data, but we might estimate an *upper bound* of 30 minutes between rupture and measurement, corresponding to 15 mV.

Liquid junction potentials (LJPs) need to be considered when a liquid-liquid interface changes after the recorded current has been ‘zeroed’ during a voltage-clamp experiment (e.g. by breaking the seal), and they are usually corrected by applying a calculated voltage offset. Typical LJP values in patch-clamp electrophysiology have been estimated as 2–12 mV (Neher, 1992). Different values are expected in different experiments, as different bath and pipette solutions are used. An appropriate correction would be expected to remove variation completely by providing the appropriate membrane voltage regardless of solutions. But failure to correct, a systematic error in the correction, or in the worst case a sign error in the correction, could lead to equal errors in both midpoints of up to 24 mV.

I_Na_ is characterised by fast time scales and large current amplitudes, both of which cause problems for membrane potential *control* in voltage-clamp experiments (Sherman et al., 1999; Lei et al., 2020; Montnach et al., 2021). In particular, a combination of cell capacitance (which increases with size) and series resistance (which depends on the quality of the seal) can cause large shifts in either midpoint. Techniques such as series resistance compensation are commonly used, but even then shifts as large as 10 mV can be incurred (Montnach et al., 2021), while under less favourable conditions shifts of 20 mV (Montnach et al., 2021) or 30 mV (Abrasheva et al., 2024) can be expected. Because the size of this effect depends on cell size and seal quality we can expect variability within experiments, and because it depends on quality control procedures and the precise technology used in the lab, we can also expect (correlated and uncorrelated) between-experiment effects, so that we classify voltage control errors as contributing to all three columns of Table 1.

Voltage step protocols vary between studies and can affect the result. For midpoints, which are steady-state properties, a major factor will be the duration of the steps intended to bring the channel into steady-state (for an example in I_Kr_ see Vandenberg et al., 2012). Different analysis methods also produce different results, with differences of up to 20 mV seen between methods (Clerx et al., 2019, again in I_Kr_). Assuming this effect depends only on the approach and not on the individual cells, but not making any assumptions about the direction of the effect, we assign it to both between-experiment columns of Table 1.

Stretch induced by deliberate pressure applied to oocytes has been shown to shift midpoints of activation by more than 10 mV (Banderali et al., 2010). If smaller amounts of pressure could be applied *accidentally*, for example by pressure from liquid flow or a badly positioned pipette, could we expect some within-experiment variability as a result? Endogenous currents are known to be present in expression systems, which can interfere with midpoint measurements (Zhang et al., 2022). Use of different cell lines, with different levels of endogenous currents, may cause between-experiment variability, while differing expression levels in each cell could cause within-experiment variability. Finally, several factors including channel glycosylation and phosphorylation regulate I_Na_ in cardiomyocytes (Marionneau and Abriel, 2015; Daimi et al., 2022). While some of these mechanisms may be highly specialised to cardiomyocytes, we might expect some forms of biological regulation even in cells non-natively expressing sodium channels, which could cause any type of variability depending on how the mechanisms themselves vary.

### Implications

The existence of substantial variability, along with the scope for error in these notoriously difficult experiments, has practical implications for experiments and data integration. First, it underscores the already well-established need for control measurements with every mutant, drug, or other moderating factor studied. And when measuring multiple mutants over a longer period of time, multiple controls also seem advised. But if we conclude that pairing is *always* necessary, then how do we interpret studies measuring the ‘canonical’ electrophysiology in a particular cell type and species (e.g. Sakakibara et al., 1992)? And does it imply that stem-cell studies always require simultaneous measurements in a healthy volunteer?

A second consequence is that data integration, combining data from different sources through numerical modelling or some other theoretical framework, can only rely on relative differences between subgroups. For example, in studies of mutants, a *shift* in a midpoint should be the preferred piece of information, while the absolute measured midpoints should be taken to be accurate only to within about 40 mV! The situation becomes more complicated if these results extrapolate to other features of I_Na_. For example, if one study investigates activation, one looks at fast inactivation, and a third at recovery, can we combine these results in a single model, or does the variability prohibit this? An emerging technology that could help address this issue is the use of short information-rich voltage-protocols, which target multiple features of ionic currents at once (Beattie et al., 2018) — although these protocols are themselves derived from preliminary modelling work on conventional protocol data. The difficulties posed by between-experiment variability for data integration are likely to also be relevant to funders, publishers, and universities, who are increasingly trying to move away from treating papers as insular results, instead trying to build strongly linked networks of reusable resources.

Thirdly, as it is possible that most of the variability is due to experimental factors that were not reported but known or easily measurable at the time, this study re-emphasises the need for greater sharing of data and meta data, already acknowledged in standards such as MICEE (Quinn et al., 2011). For example, the data set used here was created by extracting only 6 numbers per experiment from each study, while 1000s of data points were recorded originally for each *cell*. Taking advantage of modern data sharing techniques will allow future researchers to perform far more in-depth analyses. An exciting new opportunity for meta data is offered by recent USB-connected patch-clamp amplifiers, which can automatically store the applied voltage protocols, series resistance, cell capacitance, correction and compensation settings and more, all in the same file as the measured currents. This has the potential to greatly enhance what future meta analyses can do, particularly if (1) a strong data and meta data-sharing culture is established, and (2) either open-source or open-but-proprietary file formats are used (e.g. the HEKA PatchMaster format).

Finally, even when confounding variables are controlled in a single-lab multi-experiment study, a between-experiment variability of 7–11 mV remains (Tan et al., 2005; Kapplinger et al., 2015). It is a fascinating question whether this is due to as-of-yet unknown seasonal processes native to the cell, a more mundane drift in experimental conditions, or even a result of limited sample size.

## Conclusion and future directions

We reviewed 158 reported mean midpoints of activation (*µ*_*a*_) and 167 reported mean midpoints of inactivation (*µ*_*i*_), gathered from 120 publications, and found both within-experiment and between-experiment variability. Within experiments, the median standard deviation was 4.0 mV (*σ*_*a*_) or 3.6 mV (*σ*_*i*_), equivalent to 5^th^-to-95^th^ percentile ranges of 13 mV and 12 mV respectively. Between experiments, values varied over a range of 39 mV (*µ*_*a*_) or 51 mV (*µ*_*i*_), with 5^th^-to-95^th^ percentile ranges of 29 mV and 36 mV. Grouping by the known and reported confounders *α* subunit, *β*1 co-expression, and cell type did not explain this variability. In the 151 experiments providing both *µ*_*a*_ and *µ*_*i*_, we found a significant correlation with a slope almost equal to 1, hinting at some unknown factor(s) affecting both midpoints equally. While it is tempting to look for biological causes of such variability, several experimental confounders exist, which mean no such conclusions can be drawn from an analysis of the published literature. These results show that care must be taken in situations where paired experiments are not possible, or when data about different facets of channel behaviour is taken from different studies (e.g. in modelling). They also highlight the need to take full advantage of new data recording and sharing opportunities, far beyond the scope of traditional methods sections, so that future meta analyses may untangle the different possible sources of variability. We conclude that a larger than hitherto reported variability exists in the mid-points of activation and inactivation of I_Na_, and that these are highly correlated. And while the available evidence leaves room for the existence of cell-to-cell variability in the voltage-dependence of I_Na_ (with some regulatory mechanism maintaining a certain difference between the two), a simpler explanation at this point is that unreported experimental confounders give rise to the observed variability.

## References

Abrasheva VO, Kovalenko SG, Slotvitsky M, Romanova SA, Aitova AA, Frolova S, Tsvelaya V & Syunyaev RA (2024). Human sodium current voltage-dependence at physiological temperature measured by coupling a patch-clamp experiment to a mathematical model. The Journal of Physiology 602, 633–661.

Abriel H, Cabo C, Wehrens XH, Rivolta I, Motoike HK, Memmi M, Napolitano C, Priori SG & Kass RS (2001). Novel arrhythmogenic mechanism revealed by a long-QT syndrome mutation in the cardiac Na^+^ channel. Circulation Research 88, 740–745.

Ackerman MJ, Splawski I, Makielski JC, Tester DJ, Will ML, Timothy KW, Keating MT, Jones G, Chadha M, Burrow CR et al. (2004). Spectrum and prevalence of cardiac sodium channel variants among black, white, Asian, and Hispanic individuals: implications for arrhythmogenic susceptibility and Brugada/long QT syndrome genetic testing. Heart Rhythm 1, 600–607.

Banderali U, Juranka PF, Clark RB, Giles WR & Morris CE (2010). Impaired stretch modulation in potentially lethal cardiac sodium channel mutants. Channels 4, 12–21.

Beattie KA, Hill AP, Bardenet R, Cui Y, Vandenberg JI, Gavaghan DJ, de Boer TP & Mirams GR (2018). Sinusoidal voltage protocols for rapid characterization of ion channel kinetics. The Journal of Physiology 596, 1813–1828.

Beyder A, Rae JL, Bernard C, Strege PR, Sachs F & Farrugia G (2010). Mechanosensitivity of NaV1.5, a voltage-sensitive sodium channel. The Journal of physiology 588, 4969–4985.

Chadda KR, Jeevaratnam K, Lei M & Huang CLH (2017). Sodium channel biophysics, late sodium current and genetic arrhythmic syndromes. Pflügers Archiv-European Journal of Physiology 469, 629–641.

Clerx M, Beattie KA, Gavaghan DJ & Mirams GR (2019). Four ways to fit an ion channel model. Biophysical Journal 117, 2420–2437.

Clerx M, Heijman J, Collins P & Volders PGA (2018). Predicting changes to INa from missense mutations in human SCN5A. Scientific reports 8, 12797.

Daimi H, Lozano-Velasco E, Aranega A & Franco D (2022). Genomic and non-genomic regulatory mechanisms of the cardiac sodium channel in cardiac arrhythmias. International Journal of Molecular Sciences 23, 1381.

Hanck DA & Sheets MF (1992). Time-dependent changes in kinetics of Na^+^ current in single canine cardiac Purkinje cells. American Journal of Physiology. Heart and Circulatory Physiology 262, H1197–H1207.

Hedley PL, Jørgensen P, Schlamowitz S, Moolman-Smook J, Kanters JK, Corfield VA & Christiansen M (2009). The genetic basis of Brugada syndrome: a mutation update. Human Mutation 30, 1256–1266.

Hichri E, Selimi Z & Kucera JP (2020). Modeling the interactions between sodium channels provides insight into the negative dominance of certain channel mutations. Frontiers in physiology p. 1423.

Kapplinger JD, Tester DJ, Alders M, Benito B, Berthet M, Brugada J, Brugada P, Fressart V, Guerchicoff A, Harris-Kerr C et al. (2010). An international compendium of mutations in the SCN5A-encoded cardiac sodium channel in patients referred for Brugada syndrome genetic testing. Heart Rhythm 7, 33–46.

Kapplinger J, Giudicessi J, Ye D, Tester D, Callis T, Valdivia C, Makielski J, Wilde A & Ackerman M (2015). Enhanced classification of Brugada syndrome-associated and long-QT syndrome-associated genetic variants in the SCN5A-encoded Na(v)1.5 cardiac sodium channel. Circulation: Cardiovascular Genetics 8, 582–595.

Keller DI, Rougier JS, Kucera JP, Benammar N, Fressart V, Guicheney P, Madle A, Fromer M, Schläpfer J & Abriel H (2005). Brugada syndrome and fever: genetic and molecular characterization of patients carrying SCN5A mutations. Cardiovascular Research 67, 510–519.

Lei C, Clerx M, Whittaker DG, Gavaghan DJ, de Boer TP & Mirams GR (2020). Accounting for variability in ion current recordings using a mathematical model of artefacts in voltage-clamp experiments. Philosophical Transactions of the Royal Society A: Mathematical, Physical and Engineering Sciences 378, 20190348.

Makielski JC, Ye B, Valdivia CR, Pagel MD, Pu J, Tester DJ & Ackerman MJ (2003). A ubiquitous splice variant and a common polymorphism affect heterologous expression of recombinant human SCN5A heart sodium channels. Circulation Research 93, 821–828.

Marionneau C & Abriel H (2015). Regulation of the cardiac Na^+^ channel NaV1.5 by post-translational modifications. Journal of molecular and cellular cardiology 82, 36–47.

Mirams GR, Pathmanathan P, Gray RA, Challenor P & Clayton RH (2016). White paper: Uncertainty and variability in computational and mathematical models of cardiac physiology. The Journal of Physiology 594, 6833–6847.

Montnach J, Lorenzini M, Lesage A, Simon I, Nicolas S, Moreau E, Marionneau C, Baró I, De Waard M & Loussouarn G (2021). Computer modeling of whole-cell voltage-clamp analyses to delineate guidelines for good practice of manual and automated patch-clamp. Scientific reports 11, 1–16.

Moric E, Herbert E, Trusz-Gluza M, Filipecki A, Mazurek U & Wilczok T (2003). The implications of genetic mutations in the sodium channel gene (SCN5A). Europace 5, 325–334.

Nagatomo T, Fan Z, Ye B, Tonkovich G, January C, Kyle J & Makielski J (1998). Temperature dependence of early and late currents in human cardiac wild-type and long QT DeltaKPQ Na^+^ channels. American Journal of Physiology 275, H2016–H2024.

Napolitano C, Rivolta I & Priori SG (2003). Cardiac sodium channel diseases. Clinical Chemistry and Laboratory Medicine 41, 439–444.

Neher E (1992). Correction for liquid junction potentials in patch clamp experiments In Methods in enzymology, Vol. 207, pp. 123–131. Elsevier.

Olivetti G, Giordano G, Corradi D, Melissari M, Lagrasta C, Gambert & Anversa P (1995). Gender differences and aging: effects on the human heart. Journal of the American College of Cardiology 26, 1068–1079.

Page MJ, McKenzie JE, Bossuyt PM, Boutron I, Hoffmann TC, Mulrow CD, Shamseer L, Tetzlaff JM, Akl EA, Brennan SE et al. (2021). The PRISMA 2020 statement: an updated guideline for reporting systematic reviews. International journal of surgery 88, 105906.

Quinn T, Granite S, Allessie M, Antzelevitch C, Bollensdorff C, Bub G, Burton R, Cerbai E, Chen P, Delmar M et al. (2011). Minimum information about a cardiac electrophysiology experiment (MICEE): standardised reporting for model reproducibility, interoperability, and data sharing. Progress in Biophysics and Molecular Biology 107, 4–10.

Sakakibara Y, Wasserstrom JA, Furukawa T, Jia H, Arentzen CE, Hartz RS & Singer DH (1992). Characterization of the sodium current in single human atrial myocytes. Circulation Research 71, 535–546.

Schulz DJ, Goaillard JM & Marder E (2006). Variable channel expression in identified single and electrically coupled neurons in different animals. Nature Neuroscience 9, 356–362.

Sherman AJ, Shrier A & Cooper E (1999). Series resistance compensation for whole-cell patch-clamp studies using a membrane state estimator. Biophysical Journal 77, 2590–2601.

Tan BH, Valdivia CR, Rok BA, Ye B, Ruwaldt KM, Tester DJ, Ackerman MJ & Makielski JC (2005). Common human SCN5A polymorphisms have altered electrophysiology when expressed in PQ1077 splice variants. Heart Rhythm 2, 741–747.

Vandenberg JI, Perry MD, Perrin MJ, Mann SA, K. Y & Hill AP (2012). hERG K+ channels: structure, function, and clinical significance. Physiological Reviews 92, 1393–1478.

Volders PG, Sipido KR, Vos MA, Kulcsár A, Verduyn SC & Wellens HJ (1998). Cellular basis of biventricular hypertrophy and arrhythmogenesis in dogs with chronic complete atrioventricular block and acquired torsade de pointes. Circulation 98, 1136–1147.

Zhang J, Yuan H, Yao X & Chen S (2022). Endogenous ion channels expressed in human embryonic kidney (HEK-293) cells. Pflügers Archiv-European Journal of Physiology pp. 1–16.

Zimmer T & Surber R (2008). SCN5A channelopathies – an update on mutations and mechanisms. Progress in Biophysics and Molecular Biology 98, 120–136.

