## Supplemental information for "Variability in reported midpoints of (in)activation of cardiac I_Na_"

Table 1: All reviewed studies containing more than one experiment.

| Publication | Number of experiments |
|---|---|
| Kapplinger et al. (2015) | 27 |
| Tan et al. (2005) | 15 |
| Abriel et al. (2001) | 5 |
| Cheng et al. (2010) | 4 |
| Ye et al. (2003) | 2 |
| Watanabe et al. (2011) | 2 |
| Tan et al. (2006) | 2 |
| Hu et al. (2015) | 2 |
| Cheng et al. (2011) | 2 |
| Calloe et al. (2011) | 2 |
| An et al. (1998) | 2 |

Table 2: A 'histogram' view of the difference in cell counts ($|n_a - n_i|$) and how often each was encountered. The third column gives the number of occurrences as a percentage, and the final column provides the cumulative percentage (e.g. 80.8% of experiments had an $|n_a - n_i| \leq 3$).

| $\|n_a - n_i\|$ | Number of occurrences | Percentage | Cumulative percentage |
|---|---|---|---|
| 0 | 51 | 33.8% | 33.8% |
| 1 | 32 | 21.2% | 55.0% |
| 2 | 28 | 18.5% | 73.5% |
| 3 | 11 | 7.3% | 80.8% |
| 4 | 8 | 5.3% | 86.1% |
| 5 | 6 | 4.0% | 90.1% |
| 6 | 1 | 0.7% | 90.7% |
| 7 | 4 | 2.6% | 93.4% |
| 8 | 4 | 2.6% | 96.0% |
| 10 | 2 | 1.3% | 97.4% |
| 13 | 1 | 0.7% | 98.0% |
| 16 | 3 | 2.0% | 100.0% |

Table 3: All experiments reviewed in this manuscript. A structured database containing the same data is available from https://github.com/MichaelClerx/ina-midpoints.

| Study | $V_a$ | $\sigma_a$ | $n_a$ | $V_i$ | $\sigma_i$ | $n_i$ | Cell | $\alpha$ | $\beta1$ |
|---|---|---|---|---|---|---|---|---|---|
| Abe et al. (2014) | -50.5 | 5.81 | 15 | -84.1 | 5.03 | 15 | HEK | b | no |
| Abriel et al. (2000) | | | | -66.2 | 1.8 | 4 | HEK | a* | yes |
| Abriel et al. (2001) | -21.5 | 0.735 | 6 | -65.2 | 0.721 | 13 | HEK | a* | yes |
| Abriel et al. (2001) | -23.3 | 2.08 | 3 | -63.3 | 0.894 | 5 | HEK | a* | yes |
| Abriel et al. (2001) | -24.6 | 1.56 | 3 | -63.7 | 0.671 | 5 | HEK | a* | yes |
| Abriel et al. (2001) | -25.5 | 1.56 | 3 | -64.3 | 0.671 | 5 | HEK | a* | yes |
| Abriel et al. (2001) | -26 | 1.91 | 3 | -64.5 | 0.671 | 5 | HEK | a* | yes |
| Aiba et al. (2014) | -43.3 | 4.76 | 7 | -80.1 | 4.8 | 9 | HEK | ? | yes |
| Akai et al. (2000) | -44.1 | 0.9 | 9 | -80.8 | 6.3 | 9 | HEK | a* | yes |
| Amin et al. (2005) | -36.7 | 6.96 | 10 | -83.3 | 5.69 | 10 | HEK | ? | yes |
| An et al. (1998) | | | | -70.2 | 5.36 | 17 | HEK | a* | no |
| An et al. (1998) | | | | -58.7 | 4.8 | 16 | HEK | a* | yes |
| Bankston et al. (2007a) | -24.9 | 1.9 | 7 | -61.3 | 3.67 | 5 | HEK | ? | yes |
| Bankston et al. (2007b) | -24.8 | 4.69 | 13 | -71.2 | 2.7 | 9 | HEK | a* | yes |
| Baroudi et al. (2000) | | | | -101 | 5.63 | 22 | HEK | a* | no |
| Baroudi and Chahine (2000) | -47.2 | 8.15 | 23 | -93.2 | 5.18 | 21 | HEK | a* | yes |
| Baroudi et al. (2001) | -47.2 | 4.02 | 5 | -92.5 | 2.26 | 4 | HEK | a* | yes |
| Bébarová et al. (2008) | -31.8 | 4.8 | 16 | -66.6 | 3.1 | 15 | CHO | a | no |
| Beckermann et al. (2014) | -37.3 | 2.24 | 14 | -86 | 1.33 | 11 | HEK | ? | yes |
| Beyder et al. (2010) | -33 | 22 | 6 | | | | HEK | b | no |
| Beyder et al. (2014) | -58.2 | 3 | 9 | -95.5 | 3.9 | 9 | HEK | b | no |
| Calloe et al. (2011) | -34.4 | 0.566 | 8 | -71.2 | 0.9 | 9 | CHO | b | no |
| Calloe et al. (2011) | -31.4 | 1.26 | 10 | -77.7 | 1.8 | 9 | CHO | b | yes |
| Calloe et al. (2013) | -32.7 | 0.529 | 7 | -69.5 | 0.529 | 7 | CHO | b | no |
| Casini et al. (2007) | -38.6 | 3.87 | 15 | -88 | 7.57 | 13 | HEK | a* | yes |
| Chang et al. (2004) | -58.4 | 4.8 | 9 | -107 | 2.7 | 9 | HEK | ? | yes |
| Chen et al. (2016) | -45.7 | 2.62 | 14 | -80.7 | 4.5 | 12 | HEK | ? | yes |
| Cheng et al. (2010) | -32.2 | 3.2 | 16 | -79.7 | 3.71 | 17 | HEK | b | no |
| Cheng et al. (2010) | -31.1 | 3.43 | 6 | -70.6 | 3.11 | 8 | HEK | a | no |
| Cheng et al. (2010) | -34.9 | 3.6 | 9 | -72 | 2.65 | 11 | HEK | b | no |
| Cheng et al. (2010) | -34.9 | 2.01 | 5 | -72.9 | 2.65 | 7 | HEK | a | no |
| Cheng et al. (2011) | -37.6 | 3.39 | 8 | -76.1 | 4.5 | 7 | HEK | b | no |
| Cheng et al. (2011) | -39.6 | 5.59 | 5 | -74.6 | 4.23 | 7 | HEK | a | no |
| Clatot et al. (2012) | -44.3 | 6.6 | 17 | -81.7 | 1.9 | 10 | HEK | b* | no |
| Cordeiro et al. (2006) | -49.3 | 1.05 | 15 | -93 | 0.538 | 10 | HEK | b* | yes |
| Crotti et al. (2012) | -50.8 | 10.3 | 33 | -92.5 | 4.21 | 17 | HEK | ? | yes |
| Deschênes et al. (2000) | -53.6 | 4.47 | 5 | -97.4 | 2.69 | 6 | HEK | a* | yes |
| Detta et al. (2014) | -40.3 | 1.53 | 11 | | | | HEK | a* | yes |
| Ge et al. (2008) | -35.5 | 5.05 | 13 | -78.7 | 6.63 | 26 | HEK | a | yes |
| Glaaser et al. (2012) | | | | -69.1 | 9.9 | 9 | HEK | ? | no |
| Gütter et al. (2013) | -35.1 | 3.75 | 88 | -84.3 | 4.95 | 68 | HEK | a* | no |
| Gui et al. (2010a) | -34.7 | 3.36 | 23 | -81.4 | 3.43 | 24 | HEK | a* | no |

| Study | $V_a$ | $\sigma_a$ | $n_a$ | $V_i$ | $\sigma_i$ | $n_i$ | Cell | $\alpha$ | $\beta 1$ |
|---|---|---|---|---|---|---|---|---|---|
| Gui et al. (2010b) | -33.8 | 2.32 | 11 | -80.3 | 3.12 | 12 | HEK | a* | no |
| Hayashi et al. (2015) | -43.8 | 6.8 | 16 | -80 | 2.4 | 16 | CHO | ? | yes |
| Holst et al. (2009) | -30.4 | 2.24 | 14 | -82 | 4.69 | 13 | HEK | ? | no |
| Hoshi et al. (2014) | | | | -79.5 | 2.55 | 18 | HEK | a | no |
| Hsueh et al. (2009) | -42.6 | 2.56 | 10 | -84.3 | 3.79 | 10 | HEK | a | yes |
| Hu et al. (2007) | -50.8 | 1.03 | 33 | -92.5 | 0.412 | 17 | HEK | b* | yes |
| Hu et al. (2010) | | | | -92.5 | 4.21 | 17 | HEK | ? | yes |
| Hu et al. (2014) | | | | -92.5 | 3.85 | 22 | HEK | b | yes |
| Hu et al. (2015) | -41 | 3.9 | 9 | -80 | 6.97 | 19 | HEK | b | no |
| Hu et al. (2015) | -41 | 8.65 | 13 | -80 | 9.35 | 14 | HEK | a | no |
| Huang et al. (2006) | -59.9 | 2.55 | 8 | -108 | 5.09 | 8 | HEK | ? | no |
| Huang et al. (2009) | -50.3 | 6.18 | 9 | -100 | 4.02 | 9 | HEK | a* | yes |
| Itoh et al. (2005a) | -39.9 | 7.5 | 25 | | | | HEK | a* | yes |
| Itoh et al. (2005b) | -40.6 | 6.42 | 21 | | | | HEK | a* | yes |
| Itoh et al. (2007) | | | | -90.9 | 3.3 | 9 | HEK | a* | yes |
| Juang et al. (2014) | -36.3 | 0.4 | 4 | -86.4 | 1.4 | 4 | HEK | a | yes |
| Kapplinger et al. (2015) | -35.7 | 2.53 | 10 | -79.4 | 2.85 | 10 | HEK | b | no |
| Kapplinger et al. (2015) | -40 | 2.4 | 9 | -87.2 | 2.4 | 9 | HEK | b | no |
| Kapplinger et al. (2015) | -40.7 | 1.7 | 8 | -88.6 | 1.32 | 7 | HEK | a | no |
| Kapplinger et al. (2015) | -37.1 | 3.46 | 12 | -79 | 0.735 | 6 | HEK | b | no |
| Kapplinger et al. (2015) | -37.1 | 5.05 | 13 | -79.7 | 2.52 | 13 | HEK | b | no |
| Kapplinger et al. (2015) | -33.4 | 4.65 | 15 | -77.9 | 1.94 | 15 | HEK | b | no |
| Kapplinger et al. (2015) | -37 | 5.05 | 13 | -81.2 | 2.52 | 13 | HEK | b | no |
| Kapplinger et al. (2015) | -35.2 | 3.32 | 11 | -79.6 | 2.53 | 10 | HEK | b | no |
| Kapplinger et al. (2015) | -38.3 | 0.9 | 9 | -79.9 | 3.39 | 8 | HEK | a | no |
| Kapplinger et al. (2015) | -35.4 | 5.61 | 14 | -82.8 | 2.62 | 14 | HEK | b | no |
| Kapplinger et al. (2015) | -36.2 | 4.64 | 11 | -82.4 | 3.32 | 11 | HEK | a | no |
| Kapplinger et al. (2015) | -30.3 | 7.36 | 15 | -78.3 | 2.88 | 13 | HEK | b | no |
| Kapplinger et al. (2015) | -34.9 | 5.05 | 13 | -81.8 | 2.52 | 13 | HEK | b | no |
| Kapplinger et al. (2015) | -34.4 | 5.81 | 15 | -80.1 | 1.5 | 14 | HEK | b | no |
| Kapplinger et al. (2015) | -37 | 5.05 | 13 | -81.2 | 2.52 | 13 | HEK | b | no |
| Kapplinger et al. (2015) | -35.1 | 6.63 | 11 | -80.5 | 1.66 | 11 | HEK | b | no |
| Kapplinger et al. (2015) | -30.3 | 7.36 | 15 | -78.3 | 2.88 | 13 | HEK | b | no |
| Kapplinger et al. (2015) | -37.5 | 3.18 | 6 | -84.1 | 1.59 | 7 | HEK | b | no |
| Kapplinger et al. (2015) | -39.4 | 3.39 | 8 | -86.6 | 2.83 | 8 | HEK | a | no |
| Kapplinger et al. (2015) | -38.3 | 5.09 | 18 | -81.9 | 3.87 | 15 | HEK | b | no |
| Kapplinger et al. (2015) | -36.7 | 4.26 | 15 | -80.9 | 2.16 | 13 | HEK | a | no |
| Kapplinger et al. (2015) | -37.1 | 5.05 | 13 | -79.7 | 2.52 | 13 | HEK | b | no |
| Kapplinger et al. (2015) | -34.4 | 5.81 | 15 | -80.1 | 1.5 | 14 | HEK | b | no |
| Kapplinger et al. (2015) | -37.9 | 3.11 | 8 | -79.7 | 3.96 | 8 | HEK | b | no |
| Kapplinger et al. (2015) | -37.1 | 5.05 | 13 | -79.7 | 2.52 | 13 | HEK | b | no |
| Kapplinger et al. (2015) | -36.1 | 4.11 | 10 | -77.5 | 3 | 9 | HEK | b | no |
| Kapplinger et al. (2015) | -35.7 | 5.03 | 15 | -79.5 | 2.71 | 15 | HEK | b | no |
| Kato et al. (2014) | -54.4 | 8.91 | 18 | -83.8 | 9.62 | 21 | CHO | a* | yes |
| Keller et al. (2005) | -41 | 6.9 | 9 | -77.1 | 3.58 | 5 | HEK | b* | yes |

| Study | $V_a$ | $\sigma_a$ | $n_a$ | $V_i$ | $\sigma_i$ | $n_i$ | Cell | $\alpha$ | $\beta 1$ |
|---|---|---|---|---|---|---|---|---|---|
| Keller et al. (2006) | -60.1 | 4.49 | 10 | -104 | 1.81 | 8 | HEK | ? | yes |
| Li et al. (2009) | -56.6 | 4.21 | 6 | -104 | 3.87 | 6 | HEK | ? | no |
| Lin et al. (2008) | -54.6 | 1.96 | 7 | -99 | 2.46 | 8 | HEK | a* | yes |
| Liu et al. (2002) | | | | -73.3 | 6.2 | 4 | HEK | a* | yes |
| Liu et al. (2003) | -50.4 | 4.38 | 11 | -76.4 | 4.8 | 16 | HEK | ? | no |
| Liu et al. (2005) | | | | -97 | 4.2 | 9 | CHO | b | no |
| Lupoglazoff et al. (2001) | -47.2 | 7.85 | 19 | -92.5 | 3.65 | 11 | HEK | a* | yes |
| Makita et al. (2002) | -47.2 | 3.97 | 13 | -91 | 4.69 | 13 | HEK | a* | yes |
| Makita et al. (2005) | -48.1 | 3.92 | 19 | -86.6 | 3.71 | 17 | HEK | ? | yes |
| Makita et al. (2008) | -49.7 | 6.22 | 32 | -86.8 | 5.5 | 25 | HEK | ? | yes |
| Makiyama et al. (2008) | -43.6 | 3.79 | 23 | -78.1 | 4.41 | 22 | HEK | a* | yes |
| Marangoni et al. (2011) | -44 | 8 | 16 | -92 | 7.21 | 13 | HEK | ? | yes |
| Medeiros-Domingo et al. (2007) | -43.8 | 2.86 | 5 | -78.8 | 3.51 | 10 | HEK | ? | no |
| Medeiros-Domingo et al. (2009) | -38.6 | 3.49 | 15 | -76 | 5.23 | 19 | HEK | b | no |
| Mohler et al. (2004) | -41.8 | 2.65 | 7 | -68.9 | 1.16 | 15 | HEK | a* | yes |
| Mok et al. (2003) | -47.2 | 5.6 | 4 | -91.1 | 0.693 | 3 | HEK | ? | yes |
| Moreau et al. (2013) | -47.9 | 4.33 | 13 | -92 | 4.69 | 13 | HEK | a* | yes |
| Murphy et al. (2012) | -50.9 | 10.3 | 20 | -102 | 6.32 | 10 | HEK | a* | no |
| Nakajima et al. (2015) | -38.7 | 3.1 | 15 | -85.9 | 2.47 | 17 | HEK | b | yes |
| Neu et al. (2010) | -50.9 | 5.89 | 12 | -90.9 | 4.2 | 9 | HEK | a* | yes |
| Nguyen et al. (2008) | -47.8 | 1.58 | 10 | -89.4 | 2.53 | 10 | HEK | a* | yes |
| Olesen et al. (2012) | -27.9 | 6.1 | 22 | -85.6 | 4.5 | 25 | HEK | ? | no |
| Otagiri et al. (2008) | -44.4 | 4.26 | 37 | -88.3 | 4.87 | 37 | HEK | ? | yes |
| Pfahnl et al. (2007) | -50 | 1.55 | 15 | -98 | 8.52 | 15 | HEK | b* | no |
| Rivolta et al. (2001) | -23.3 | 2.24 | 5 | -62.8 | 3.12 | 12 | HEK | ? | yes |
| Rossenbacker et al. (2004) | -24.1 | 0.894 | 5 | -70.9 | 1.4 | 4 | HEK | ? | yes |
| Ruan et al. (2007) | -23.2 | 1.92 | 5 | -62.5 | 2.15 | 10 | HEK | ? | yes |
| Ruan et al. (2010) | -23.1 | 1.77 | 9 | -67.7 | 3.01 | 12 | HEK | ? | yes |
| Saber et al. (2015) | -24 | 4.9 | 6 | -66 | 4.9 | 6 | HEK | ? | yes |
| Samani et al. (2009) | -36 | 5.03 | 7 | -89.9 | 5.4 | 9 | HEK | b | yes |
| Sarhan et al. (2009) | | | | -109 | 1.85 | 7 | HEK | a | no |
| Shinlapawittayatorn et al. (2011b) | | | | -91.9 | 4.5 | 7 | HEK | a | no |
| Shinlapawittayatorn et al. (2011a) | | | | -91.2 | 2.77 | 12 | HEK | a | no |
| Shirai et al. (2002) | -49.9 | 2.38 | 7 | -94.9 | 6.37 | 6 | HEK | a* | no |
| Shuraih et al. (2007) | -43.4 | 0.794 | 7 | -90.7 | 0.265 | 7 | HEK | b | yes |
| Shy et al. (2014) | -29.6 | 3.68 | 8 | -76.9 | 6.96 | 10 | HEK | ? | no |
| Smits et al. (2005a) | -42.6 | 4.2 | 9 | -89.4 | 3.6 | 9 | HEK | a* | yes |
| Smits et al. (2005b) | -43.7 | 9 | 9 | -98.8 | 7.57 | 13 | HEK | a* | yes |
| Sottas et al. (2013) | -29.8 | 1.99 | 11 | -72.2 | 1.66 | 11 | HEK | ? | yes |
| Splawski et al. (2002) | -26.6 | 3.68 | 8 | | | | HEK | ? | no |
| Surber et al. (2008) | -42.2 | 3.37 | 14 | -79.4 | 3.43 | 6 | HEK | a* | no |
| Swan et al. (2014) | -28 | 3.39 | 8 | -75.8 | 4.9 | 6 | HEK | b | yes |
| Tan et al. (2001) | -48.6 | 3.17 | 7 | -92 | 4.5 | 7 | HEK | ? | yes |
| Tan et al. (2002) | -40.3 | 2.88 | 13 | -93.5 | 4.33 | 13 | HEK | a* | no |
| Tan et al. (2005) | -39 | 4.9 | 6 | -75 | 5.66 | 8 | HEK | b | no |

| Study | $V_a$ | $\sigma_a$ | $n_a$ | $V_i$ | $\sigma_i$ | $n_i$ | Cell | $\alpha$ | $\beta1$ |
|---|---|---|---|---|---|---|---|---|---|
| Tan et al. (2005) | -38 | 4.9 | 6 | -75 | 6 | 9 | HEK | b | no |
| Tan et al. (2005) | -42 | 2.91 | 5 | -81 | 4.2 | 9 | HEK | b | no |
| Tan et al. (2005) | -41 | 3.39 | 8 | -79 | 5.31 | 11 | HEK | b | no |
| Tan et al. (2005) | -42 | 2.55 | 8 | -79 | 4.2 | 9 | HEK | b | no |
| Tan et al. (2005) | -40 | 9.55 | 6 | -78 | 7.83 | 5 | HEK | b | no |
| Tan et al. (2005) | -40 | 1.39 | 3 | -81 | 4.68 | 3 | HEK | b | no |
| Tan et al. (2005) | -39 | 4 | 4 | -75 | 5.66 | 8 | HEK | a | no |
| Tan et al. (2005) | -39 | 4 | 4 | -78 | 2.83 | 8 | HEK | a | no |
| Tan et al. (2005) | -42 | 3.96 | 8 | -82 | 4.85 | 12 | HEK | a | no |
| Tan et al. (2005) | -40 | 8.49 | 18 | -79 | 8.49 | 18 | HEK | a | no |
| Tan et al. (2005) | -43 | 2 | 4 | -82 | 6.2 | 4 | HEK | a | no |
| Tan et al. (2005) | -42 | 3.39 | 8 | -82 | 3.6 | 9 | HEK | a | no |
| Tan et al. (2005) | -40 | 5.66 | 8 | -80 | 4.8 | 9 | HEK | a | no |
| Tan et al. (2005) | -41 | 1.8 | 4 | -80 | 1.2 | 4 | HEK | a | no |
| Tan et al. (2006) | -46.9 | 3.39 | 8 | -81.8 | 3.68 | 8 | HEK | b | no |
| Tan et al. (2006) | -44.1 | 5.06 | 10 | -80 | 4.74 | 10 | HEK | a | no |
| Tarradas et al. (2013) | -32 | 1.27 | 18 | -84.9 | 2.85 | 10 | HEK | a* | no |
| Tester et al. (2010) | -42 | 4 | 4 | -72 | 2.24 | 5 | HEK | b | no |
| Tsurugi et al. (2009) | -39.6 | 3.39 | 8 | -88 | 2.55 | 8 | HEK | ? | no |
| Valdivia et al. (2004) | -42 | 7.75 | 15 | -84.3 | 4.47 | 20 | HEK | b | no |
| Vatta et al. (2002) | | | | -89.5 | 0.49 | 6 | HEK | ? | no |
| Viswanathan et al. (2003) | -40.7 | 4.64 | 11 | -85 | 3.98 | 11 | HEK | a* | yes |
| Wang et al. (1996) | -43.2 | 6.85 | 13 | -99.6 | 2.92 | 11 | HEK | a* | no |
| Wang et al. (2002) | -47.7 | 4 | 16 | -101 | 6.1 | 19 | HEK | a* | yes |
| Wang et al. (2007) | -44.3 | 2.24 | 14 | -89.3 | 4.4 | 16 | HEK | a* | yes |
| Wang et al. (2007) | -46 | 6.93 | 5 | | | | HEK | a* | yes |
| Wang et al. (2008) | -44.3 | 2.24 | 14 | -89.3 | 4.4 | 16 | HEK | a* | yes |
| Wang et al. (2011) | | | | -93.9 | 2.65 | 11 | HEK | ? | no |
| Wang et al. (2015) | -40.9 | 0.63 | 9 | -72.7 | 2.49 | 7 | HEK | a* | no |
| Wang et al. (2016) | -44.5 | 4.8 | 36 | -93.5 | 4.08 | 34 | HEK | ? | yes |
| Watanabe et al. (2011) | -35.4 | 3 | 25 | -84.5 | 4.9 | 24 | CHO | a | no |
| Watanabe et al. (2011) | -47.7 | 4.4 | 16 | -89.4 | 3.05 | 19 | HEK | a | no |
| Wedekind et al. (2001) | -42.8 | 7.67 | 7 | -98.1 | 5.03 | 7 | HEK | a* | yes |
| Wehrens et al. (2003) | -29.8 | 1.13 | 8 | -64 | 2.26 | 8 | HEK | ? | yes |
| Winkel et al. (2012) | -26 | 10.3 | 17 | -84.6 | 7.2 | 16 | HEK | ? | no |
| Yang et al. (2002) | -54.3 | 9.26 | 7 | -98.3 | 0.794 | 7 | HEK | a* | yes |
| Ye et al. (2003) | -44 | 15.8 | 10 | -95 | 14.4 | 9 | HEK | a* | no |
| Ye et al. (2003) | -40 | 18.5 | 7 | -86 | 15.3 | 7 | HEK | b* | no |
| Yokoi et al. (2005) | -49.8 | 3.68 | 8 | -88.6 | 3 | 9 | HEK | ? | no |
| Young and Caldwell (2005) | -32.7 | 5.81 | 20 | -66 | 8.94 | 20 | CHO | a* | no |
| Zeng et al. (2013) | -34.5 | 4.24 | 8 | -81.1 | 4.69 | 13 | HEK | a | yes |
| Zhang et al. (2015) | -28.1 | 5.03 | 15 | | | | HEK | ? | no |

### References

Abe K, Machida T, Sumitomo N, Yamamoto H, Ohkubo K, Watanabe I, Makiyama T, Fukae S, Kohno M, Harrell DT et al. (2014). Sodium channelopathy underlying familial sick sinus syndrome with early onset and predominantly male characteristics. *Circulation: Arrhythmia and Electrophysiology* **7**, 511–517.

Abriel H, Cabo C, Wehrens XH, Rivolta I, Motoike HK, Memmi M, Napolitano C, Priori SG & Kass RS (2001). Novel arrhythmogenic mechanism revealed by a long-QT syndrome mutation in the cardiac Na+ channel. *Circulation Research* **88**, 740–745.

Abriel H, Wehrens X, Benhorin J, Kerem B & Kass R (2000). Molecular pharmacology of the sodium channel mutation D1790G linked to the long-QT syndrome. *Circulation* **102**, 921–925.

Aiba T, Farinelli F, Kostecki G, Hesketh GG, Edwards D, Biswas S, Tung L & Tomaselli GF (2014). A mutation causing Brugada syndrome identifies a mechanism for altered autonomic and oxidant regulation of cardiac sodium currents. *Circulation: Cardiovascular Genetics* **7**, 249–256.

Akai J, Makita N, Sakurada H, Shirai N, Ueda K, Kitabatake A, Nakazawa K, Kimura A & Hiraoka M (2000). A novel SCN5A mutation associated with idiopathic ventricular fibrillation without typical ECG findings of Brugada syndrome. *FEBS Letters* **479**, 29–34.

Amin A, Verkerk A, Bhuiyan Z, Wilde A & Tan H (2005). Novel Brugada syndrome-causing mutation in ion-conducting pore of cardiac Na+ channel does not affect ion selectivity properties. *Acta Physiologica Scandinavica* **185**, 291–301.

An R, Wang X, Kerem B, Benhorin J, Medina A, Goldmit M & Kass R (1998). Novel LQT-3 mutation affects Na+ channel activity through interactions between $\alpha$-and $\beta$1-subunits. *Circulation Research* **83**, 141–146.

Bankston JR, Sampson KJ, Kateriya S, Glaaser IW, Malito DL, Chung WK & Kass RS (2007a). A novel LQT-3 mutation disrupts an inactivation gate complex with distinct rate-dependent phenotypic consequences. *Channels* **1**, 273–280.

Bankston JR, Yue M, Chung W, Spyres M, Pass RH, Silver E, Sampson KJ & Kass RS (2007b). A novel and lethal de novo LQT-3 mutation in a newborn with distinct molecular pharmacology and therapeutic response. *PLOS ONE* **2**, e1258.

Baroudi G, Carbonneau E, Pouliot V & Chahine M (2000). SCN5A mutation (T1620M) causing Brugada syndrome exhibits different phenotypes when expressed in Xenopus oocytes and mammalian cells. *FEBS Letters* **467**, 12–16.

Baroudi G & Chahine M (2000). Biophysical phenotypes of SCN5A mutations causing long QT and Brugada syndromes. *FEBS Letters* **487**, 224–228.

Baroudi G, Pouliot V, Denjoy I, Guicheney P, Shrier A & Chahine M (2001). Novel mechanism for Brugada syndrome defective surface localization of an SCN5A mutant (R1432G). *Circulation Research* **88**, e78–e83.

Bébarová M, O'Hara T, Geelen JL, Jongbloed RJ, Timmermans C, Arens YH, Rodriguez LM, Rudy Y & Volders PG (2008). Subepicardial phase 0 block and discontinuous transmural conduction underlie right precordial ST-segment elevation by a SCN5A loss-of-function mutation. *American Journal of Physiology – Heart and Circulatory Physiology* **295**, H48–H58.

Beckermann TM, McLeod K, Murday V, Potet F & George AL (2014). Novel SCN5A mutation in amiodarone-responsive multifocal ventricular ectopy-associated cardiomyopathy. *Heart Rhythm* **11**, 1446–1453.

Beyder A, Mazzone A, Strege PR, Tester DJ, Saito YA, Bernard CE, Enders FT, Ek WE, Schmidt PT, Dlugosz A et al. (2014). Loss-of-function of the voltage-gated sodium channel NaV1.5 (channelopathies) in patients with irritable bowel syndrome. *Gastroenterology* **146**, 1659–1668.

Beyder A, Rae JL, Bernard C, Strege PR, Sachs F & Farrugia G (2010). Mechanosensitivity of NaV1.5, a voltage-sensitive sodium channel. *The Journal of Physiology* **588**, 4969–4985.

Calloe K, Refaat MM, Grubb S, Wojciak J, Campagna J, Thomsen NM, Nussbaum RL, Scheinman MM & Schmitt N (2013). Characterization and mechanisms of action of novel NaV1.5 channel mutations associated with Brugada syndrome. *Circulation: Arrhythmia and Electrophysiology* **6**, 177–184.

Calloe K, Schmitt N, Grubb S, Pfeiffer R, David JP, Kanter R, Cordeiro JM & Antzelevitch C (2011). Multiple arrhythmic syndromes in a newborn, owing to a novel mutation in SCN5A. *Canadian Journal of Physiology and Pharmacology* **89**, 723–736.

Casini S, Tan HL, Bhuiyan ZA, Bezzina CR, Barnett P, Cerbai E, Mugelli A, Wilde AA & Veldkamp MW (2007). Characterization of a novel SCN5A mutation associated with Brugada syndrome reveals involvement of DIIIS4–S5 linker in slow inactivation. *Cardiovascular Research* **76**, 418–429.

Chang CC, Acharfi S, Wu MH, Chiang FT, Wang JK, Sung TC & Chahine M (2004). A novel SCN5A mutation manifests as a malignant form of long QT syndrome with perinatal onset of tachycardia/bradycardia. *Cardiovascular Research* **64**, 268–278.

Chen J, Makiyama T, Wuriyanghai Y, Ohno S, Sasaki K, Hayano M, Harita T, Nishiuchi S, Yamamoto Y, Ueyama T et al. (2016). Cardiac sodium channel mutation associated with epinephrine-induced QT prolongation and sinus node dysfunction. *Heart Rhythm* **13**, 289–298.

Cheng J, Morales A, Siegfried JD, Li D, Norton N, Song J, Gonzalez-Quintana J, Makielski JC & Hershberger RE (2010). SCN5A rare variants in familial dilated cardiomyopathy decrease peak sodium current depending on the common polymorphism H558R and common splice variant Q1077del. *Clinical and Translational Science* **3**, 287–294.

Cheng J, Tester DJ, Tan BH, Valdivia CR, Kroboth S, Ye B, January CT, Ackerman MJ & Makielski JC (2011). The common African American polymorphism SCN5A-S1103Y interacts with mutation SCN5A-R680H to increase late Na current. *Physiological Genomics* **43**, 461–466.

Clatot J, Ziyadeh-Isleem A, Maugenre S, Denjoy I, Liu H, Dilanian G, Hatem SN, Deschênes I, Coulombe A, Guicheney P et al. (2012). Dominant-negative effect of SCN5A N-terminal mutations through the interaction of NaV1.5 $\alpha$-subunits. *Cardiovascular Research* **96**, 53–63.

Cordeiro JM, Barajas-Martinez H, Hong K, Burashnikov E, Pfeiffer R, Orsino AM, Wu YS, Hu D, Brugada J, Brugada P et al. (2006). Compound heterozygous mutations P336L and I1660V in the human cardiac sodium channel associated with the Brugada syndrome. *Circulation* **114**, 2026–2033.

Crotti L, Hu D, Barajas-Martinez H, De Ferrari GM, Oliva A, Insolia R, Pollevick GD, Dagradi F, Guerchicoff A, Greco F et al. (2012). Torsades de pointes following acute myocardial infarction: evidence for a deadly link with a common genetic variant. *Heart Rhythm* **9**, 1104–1112.

Deschênes I, Baroudi G, Berthet M, Barde I, Chalvidan T, Denjoy I, Guicheney P & Chahine M (2000). Electrophysiological characterization of SCN5A mutations causing long QT (E1784K) and Brugada (R1512W and R1432G) syndromes. *Cardiovascular Research* **46**, 55–65.

Detta N, Frisso G, Limongelli G, Marzullo M, Calabrò R & Salvatore F (2014). Genetic analysis in a family affected by sick sinus syndrome may reduce the sudden death risk in a young aspiring competitive athlete. *International Journal of Cardiology* **170**, e63–e65.

Ge J, Sun A, Paajanen V, Wang S, Su C, Yang Z, Li Y, Wang S, Jia J, Wang K et al. (2008). Molecular and clinical characterization of a novel SCN5A mutation associated with atrioventricular block and dilated cardiomyopathy. *Circulation: Arrhythmia and Electrophysiology* **1**, 83–92.

Glaaser IW, Osteen JD, Puckerin A, Sampson KJ, Jin X & Kass RS (2012). Perturbation of sodium channel structure by an inherited long QT syndrome mutation. *Nature Communications* **3**, 706.

Gui J, Wang T, Jones RP, Trump D, Zimmer T & Lei M (2010a). Multiple loss-of-function mechanisms contribute to SCN5A-related familial sick sinus syndrome. *PLOS ONE* **5**, e10985.

Gui J, Wang T, Trump D, Zimmer T & Lei M (2010b). Mutation-specific effects of polymorphism H558R in SCN5A-related sick sinus syndrome. *Journal of Cardiovascular Electrophysiology* **21**, 564–573.

Gütter C, Benndorf K & Zimmer T (2013). Characterization of N-terminally mutated cardiac Na+ channels associated with long QT syndrome 3 and Brugada syndrome. *Sudden arrhythmic death: from basic science to clinical practice* p. 41.

Hayashi K, Konno T, Tada H, Tani S, Liu L, Fujino N, Nohara A, Hodatsu A, Tsuda T, Tanaka Y et al. (2015). Functional characterization of rare variants implicated in susceptibility to lone atrial fibrillation. *Circulation: Arrhythmia and Electrophysiology* **8**, 1095–1104.

Holst AG, Calloe K, Jespersen T, Cedergreen P, Winkel BG, Jensen HK, Leren TP, Haunso S, Svendsen JH & Tfelt-Hansen J (2009). A novel SCN5A mutation in a patient with coexistence of Brugada syndrome traits and ischaemic heart disease. *Case Reports in Medicine* **2009**.

Hoshi M, Du XX, Shinlapawittayatorn K, Liu H, Chai S, Wan X, Ficker E & Deschênes I (2014). Brugada syndrome disease phenotype explained in apparently benign sodium channel mutations. *Circulation: Cardiovascular Genetics* **7**, 123.

Hsueh CH, Chen WP, Lin JL, Tsai CT, Liu YB, Juang JM, Tsao HM, Su MJ & Lai LP (2009). Distinct functional defect of three novel Brugada syndrome related cardiac sodium channel mutations. *Journal of Biomedical Science* **16**, 1.

Hu D, Barajas-Martinez H, Nesterenko VV, Pfeiffer R, Guerchicoff A, Cordeiro JM, Curtis AB, Pollevick GD, Wu Y, Burashnikov E et al. (2010). Dual variation in SCN5A and CACNB2b underlies the development of cardiac conduction disease without Brugada syndrome. *Pacing and Clinical Electrophysiology* **33**, 274–285.

Hu D, Barajas-Martínez H, Terzic A, Park S, Pfeiffer R, Burashnikov E, Wu Y, Borggrefe M, Veltmann C, Schimpf R et al. (2014). ABCC9 is a novel Brugada and early repolarization syndrome susceptibility gene. *International Journal of Cardiology* **171**, 431–442.

Hu D, Viskin S, Oliva A, Carrier T, Cordeiro JM, Barajas-Martinez H, Wu Y, Burashnikov E, Sicouri S, Brugada R et al. (2007). Novel mutation in the SCN5A gene associated with arrhythmic storm development during acute myocardial infarction. *Heart Rhythm* **4**, 1072–1080.

Hu RM, Tan BH, Tester DJ, Song C, He Y, Dovat S, Peterson BZ, Ackerman MJ & Makielski JC (2015). Arrhythmogenic biophysical phenotype for SCN5A mutation S1787N depends upon splice variant background and intracellular acidosis. *PLOS ONE* **10**, e0124921.

Huang H, Millat G, Rodriguez-Lafrasse C, Rousson R, Kugener B, Chevalier P & Chahine M (2009). Biophysical characterization of a new SCN5A mutation S1333Y in a SIDS infant linked to long QT syndrome. *FEBS Letters* **583**, 890–896.

Huang H, Zhao J, Barrane FZ, Champagne J & Chahine M (2006). NaV1.5/R1193Q polymorphism is associated with both long QT and Brugada syndromes. *Canadian Journal of Cardiology* **22**, 309–313.

Itoh H, Shimizu M, Mabuchi H & Imoto K (2005a). Clinical and electrophysiological characteristics of Brugada syndrome caused by a missense mutation in the S5-pore site of SCN5A. *Journal of Cardiovascular Electrophysiology* **16**, 378–383.

Itoh H, Shimizu M, Takata S, Mabuchi H & Imoto K (2005b). A novel missense mutation in the SCN5A gene associated with Brugada syndrome bidirectionally affecting blocking actions of antiarrhythmic drugs. *Journal of Cardiovascular Electrophysiology* **16**, 486–493.

Itoh H, Tsuji K, Sakaguchi T, Nagaoka I, Oka Y, Nakazawa Y, Yao T, Jo H, Ashihara T, Ito M et al. (2007). A paradoxical effect of lidocaine for the N406S mutation of SCN5A associated with Brugada syndrome. *International Journal of Cardiology* **121**, 239–248.

Juang JMJ, Lu TP, Lai LC, Hsueh CH, Liu YB, Tsai CT, Lin LY, Yu CC, Hwang JJ, Chiang FT et al. (2014). Utilizing multiple in silico analyses to identify putative causal SCN5A variants in Brugada syndrome. *Scientific Reports* **4**, 3850.

Kapplinger J, Giudicessi J, Ye D, Tester D, Callis T, Valdivia C, Makielski J, Wilde A & Ackerman M (2015). Enhanced classification of Brugada syndrome-associated and long-QT syndrome-associated genetic variants in the SCN5A-encoded Na(v)1.5 cardiac sodium channel. *Circulation: Cardiovascular Genetics* **8**, 582–595.

Kato K, Makiyama T, Wu J, Ding WG, Kimura H, Naiki N, Ohno S, Itoh H, Nakanishi T, Matsuura H et al. (2014). Cardiac channelopathies associated with infantile fatal ventricular arrhythmias: from the cradle to the bench. *Journal of Cardiovascular Electrophysiology* **25**, 66–73.

Keller DI, Huang H, Zhao J, Frank R, Suarez V, Delacrétaz E, Brink M, Osswald S, Schwick N & Chahine M (2006). A novel SCN5A mutation, F1344S, identified in a patient with Brugada syndrome and fever-induced ventricular fibrillation. *Cardiovascular Research* **70**, 521–529.

Keller DI, Rougier JS, Kucera JP, Benammar N, Fressart V, Guicheney P, Madle A, Fromer M, Schläpfer J & Abriel H (2005). Brugada syndrome and fever: genetic and molecular characterization of patients carrying SCN5A mutations. *Cardiovascular Research* **67**, 510–519.

Li Q, Huang H, Liu G, Lam K, Rutberg J, Green MS, Birnie DH, Lemery R, Chahine M & Gollob MH (2009). Gain-of-function mutation of NaV1.5 in atrial fibrillation enhances cellular excitability and lowers the threshold for action potential firing. *Biochemical and Biophysical Research Communications* **380**, 132–137.

Lin MT, Wu MH, Chang CC, Chiu SN, Thériault O, Huang H, Christé G, Ficker E & Chahine M (2008). In utero onset of long QT syndrome with atrioventricular block and spontaneous or lidocaine-induced ventricular tachycardia: compound effects of hERG pore region mutation and SCN5A N-terminus variant. *Heart Rhythm* **5**, 1567–1574.

Liu Cj, Dib-Hajj SD, Renganathan M, Cummins TR & Waxman SG (2003). Modulation of the cardiac sodium channel NaV1.5 by fibroblast growth factor homologous factor 1B. *Journal of Biological Chemistry* **278**, 1029–1036.

Liu H, Tateyama M, Clancy CE, Abriel H & Kass RS (2002). Channel openings are necessary but not sufficient for use-dependent block of cardiac Na+ channels by flecainide evidence from the analysis of disease-linked mutations. *The Journal of General Physiology* **120**, 39–51.

Liu K, Yang T, Viswanathan PC & Roden DM (2005). New mechanism contributing to drug-induced arrhythmia rescue of a misprocessed LQT3 mutant. *Circulation* **112**, 3239–3246.

Lupoglazoff J, Cheav T, Baroudi G, Berthet M, Denjoy I, Cauchemez B, Extramiana F, Chahine M & Guicheney P (2001). Homozygous SCN5A mutation in long-QT syndrome with functional two-to-one atrioventricular block. *Circulation Research* **89**, e16–e21.

Makita N, Behr E, Shimizu W, Horie M, Sunami A, Crotti L, Schulze-Bahr E, Fukuhara S, Mochizuki N, Makiyama T et al. (2008). The E1784K mutation in SCN5A is associated with mixed clinical phenotype of type 3 long QT syndrome. *The Journal of Clinical Investigation* **118**, 2219–2229.

Makita N, Horie M, Nakamura T, Ai T, Sasaki K, Yokoi H, Sakurai M, Sakuma I, Otani H, Sawa H et al. (2002). Drug-induced long-QT syndrome associated with a subclinical SCN5A mutation. *Circulation* **106**, 1269–1274.

Makita N, Sasaki K, Groenewegen WA, Yokota T, Yokoshiki H, Murakami T & Tsutsui H (2005). Congenital atrial standstill associated with coinheritance of a novel SCN5A mutation and connexin 40 polymorphisms. *Heart Rhythm* **2**, 1128–1134.

Makiyama T, Akao M, Shizuta S, Doi T, Nishiyama K, Oka Y, Ohno S, Nishio Y, Tsuji K, Itoh H et al. (2008). A novel SCN5A gain-of-function mutation M1875T associated with familial atrial fibrillation. *Journal of the American College of Cardiology* **52**, 1326–1334.

Marangoni S, Di Resta C, Rocchetti M, Barile L, Rizzetto R, Summa A, Severi S, Sommariva E, Pappone C, Ferrari M et al. (2011). A Brugada syndrome mutation (p.S216L) and its modulation by p.H558R polymorphism: standard and dynamic characterization. *Cardiovascular Research* **91**, 606–616.

Medeiros-Domingo A, Kaku T, Tester DJ, Iturralde-Torres P, Itty A, Ye B, Valdivia C, Ueda K, Canizales-Quinteros S, Tusié-Luna MT et al. (2007). SCN4B-encoded sodium channel $\beta 4$ subunit in congenital long-QT syndrome. *Circulation* **116**, 134–142.

Medeiros-Domingo A, Tan BH, Iturralde-Torres P, Tester DJ, Tusié-Luna T, Makielski JC & Ackerman MJ (2009). Unique mixed phenotype and unexpected functional effect revealed by novel compound heterozygosity mutations involving SCN5A. *Heart Rhythm* **6**, 1170–1175.

Mohler PJ, Rivolta I, Napolitano C, LeMaillet G, Lambert S, Priori SG & Bennett V (2004). NaV1.5 E1053K mutation causing Brugada syndrome blocks binding to ankyrin-G and expression of NaV1.5 on the surface of cardiomyocytes. *Proceedings of the National Academy of Sciences of the United States of America* **101**, 17533–17538.

Mok NS, Priori SG, Napolitano C, Chan NY, Chahine M & Baroudi G (2003). A newly characterized SCN5A mutation underlying Brugada syndrome unmasked by hyperthermia. *Journal of Cardiovascular Electrophysiology* **14**, 407–411.

Moreau A, Krahn AD, Gosselin-Badaroudine P, Klein GJ, Christé G, Vincent Y, Boutjdir M & Chahine M (2013). Sodium overload due to a persistent current that attenuates the arrhythmogenic potential of a novel LQT3 mutation. *Frontiers in Pharmacology* **4**, 126.

Murphy LL, Moon-Grady AJ, Cuneo BF, Wakai RT, Yu S, Kunic JD, Benson DW & George AL (2012). Developmentally regulated SCN5A splice variant potentiates dysfunction of a novel mutation associated with severe fetal arrhythmia. *Heart Rhythm* **9**, 590–597.

Nakajima T, Kaneko Y, Saito A, Ota M, Iijima T & Kurabayashi M (2015). Enhanced fast-inactivated state stability of cardiac sodium channels by a novel voltage sensor SCN5A mutation, R1632C, as a cause of atypical Brugada syndrome. *Heart Rhythm* **12**, 2296–2304.

Neu A, Eiselt M, Paul M, Sauter K, Stallmeyer B, Isbrandt D & Schulze-Bahr E (2010). A homozygous SCN5A mutation in a severe, recessive type of cardiac conduction disease. *Human Mutation* **31**, E1609–E1621.

Nguyen TP, Wang DW, Rhodes TH & George AL (2008). Divergent biophysical defects caused by mutant sodium channels in dilated cardiomyopathy with arrhythmia. *Circulation Research* **102**, 364–371.

Olesen MS, Yuan L, Liang B, Holst AG, Nielsen N, Nielsen JB, Hedley PL, Christiansen M, Olesen SP, Haunsø S et al. (2012). High prevalence of long QT syndrome associated SCN5A variants in patients with early-onset lone atrial fibrillation. *Circulation: Cardiovascular Genetics* **5**, 450.

Otagiri T, Kijima K, Osawa M, Ishii K, Makita N, Matoba R, Umetsu K & Hayasaka K (2008). Cardiac ion channel gene mutations in sudden infant death syndrome. *Pediatric Research* **64**, 482–487.

Pfahnl AE, Viswanathan PC, Weiss R, Shang LL, Sanyal S, Shusterman V, Kornblit C, London B & Dudley SC (2007). A sodium channel pore mutation causing Brugada syndrome. *Heart Rhythm* **4**, 46–53.

Rivolta I, Abriel H, Tateyama M, Liu H, Memmi M, Vardas P, Napolitano C, Priori SG & Kass RS (2001). Inherited Brugada and long QT-3 syndrome mutations of a single residue of the cardiac sodium channel confer distinct channel and clinical phenotypes. *Journal of Biological Chemistry* **276**, 30623–30630.

Rossenbacker T, Carroll SJ, Liu H, Kuipéri C, de Ravel TJ, Devriendt K, Carmeliet P, Kass RS & Heidbüchel H (2004). Novel pore mutation in SCN5A manifests as a spectrum of phenotypes ranging from atrial flutter, conduction disease, and Brugada syndrome to sudden cardiac death. *Heart Rhythm* **1**, 610–615.

Ruan Y, Denegri M, Liu N, Bachetti T, Seregni M, Morotti S, Severi S, Napolitano C & Priori SG (2010). Trafficking defects and gating abnormalities of a novel SCN5A mutation question gene-specific therapy in long QT syndrome type 3. *Circulation Research* **106**, 1374–1383.

Ruan Y, Liu N, Bloise R, Napolitano C & Priori SG (2007). Gating properties of SCN5A mutations and the response to mexiletine in long-QT syndrome type 3 patients. *Circulation* **116**, 1137–1144.

Saber S, Amarouch MY, Fazelifar AF, Haghjoo M, Emkanjoo Z, Alizadeh A, Houshmand M, Gavrilenko AV, Abriel H & Zaklyazminskaya EV (2015). Complex genetic background in a large family with Brugada syndrome. *Physiological Reports* **3**, e12256.

Samani K, Wu G, Ai T, Shuraih M, Mathuria NS, Li Z, Sohma Y, Purevjav E, Xi Y, Towbin JA et al. (2009). A novel SCN5A mutation V1340I in Brugada syndrome augmenting arrhythmias during febrile illness. *Heart Rhythm* **6**, 1318–1326.

Sarhan MF, Van Petegem F & Ahern CA (2009). A double tyrosine motif in the cardiac sodium channel domain III-IV linker couples calcium-dependent calmodulin binding to inactivation gating. *Journal of Biological Chemistry* **284**, 33265–33274.

Shinlapawittayatorn K, Du XX, Liu H, Ficker E, Kaufman ES & Deschênes I (2011a). A common SCN5A polymorphism modulates the biophysical defects of SCN5A mutations. *Heart Rhythm* **8**, 455–462.

Shinlapawittayatorn K, Dudash LA, Du XX, Heller L, Poelzing S, Ficker E & Deschênes I (2011b). A novel strategy using cardiac sodium channel polymorphic fragments to rescue trafficking-deficient SCN5A mutations. *Circulation: Cardiovascular Genetics* **4**, 500–509.

Shirai N, Makita N, Sasaki K, Yokoi H, Sakuma I, Sakurada H, Akai J, Kimura A, Hiraoka M & Kitabatake A (2002). A mutant cardiac sodium channel with multiple biophysical defects associated with overlapping clinical features of Brugada syndrome and cardiac conduction disease. *Cardiovascular Research* **53**, 348–354.

Shuraih M, Ai T, Vatta M, Sohma Y, Merkle EM, Taylor E, Li Z, Xi Y, Razavi M, Towbin JA et al. (2007). A common SCN5A variant alters the responsiveness of human sodium channels to class I antiarrhythmic agents. *Journal of Cardiovascular Electrophysiology* **18**, 434–440.

Shy D, Gillet L, Ogrodnik J, Albesa M, Verkerk AO, Wolswinkel R, Rougier JS, Barc J, Essers MC, Syam N et al. (2014). PDZ domain–binding motif regulates cardiomyocyte compartment-specific NaV1.5 channel expression and function. *Circulation* **130**, 147–160.

Smits JP, Koopmann TT, Wilders R, Veldkamp MW, Opthof T, Bhuiyan ZA, Mannens MM, Balser JR, Tan HL, Bezzina CR et al. (2005a). A mutation in the human cardiac sodium channel (E161K) contributes to sick sinus syndrome, conduction disease and Brugada syndrome in two families. *Journal of Molecular and Cellular Cardiology* **38**, 969–981.

Smits JP, Veldkamp MW, Bezzina CR, Bhuiyan ZA, Wedekind H, Schulze-Bahr E & Wilde AA (2005b). Substitution of a conserved alanine in the domain IIIS4–S5 linker of the cardiac sodium channel causes long QT syndrome. *Cardiovascular Research* **67**, 459–466.

Sottas V, Rougier JS, Jousset F, Kucera JP, Shestak A, Makarov LM, Zaklyazminskaya EV & Abriel H (2013). Characterization of 2 genetic variants of NaV1.5-Arginine 689 found in patients with cardiac arrhythmias. *Journal of Cardiovascular Electrophysiology* **24**, 1037–1046.

Splawski I, Timothy KW, Tateyama M, Clancy CE, Malhotra A, Beggs AH, Cappuccio FP, Sagnella GA, Kass RS & Keating MT (2002). Variant of SCN5A sodium channel implicated in risk of cardiac arrhythmia. *Science* **297**, 1333–1336.

Surber R, Hensellek S, Prochnau D, Werner GS, Benndorf K, Figulla HR & Zimmer T (2008). Combination of cardiac conduction disease and long QT syndrome caused by mutation T1620K in the cardiac sodium channel. *Cardiovascular Research* **77**, 740–748.

Swan H, Amarouch MY, Leinonen J, Marjamaa A, Kucera JP, Laitinen-Forsblom PJ, Lahtinen AM, Palotie A, Kontula K, Toivonen L et al. (2014). A gain-of-function mutation of the SCN5A gene causes exercise-induced polymorphic ventricular arrhythmias. *Circulation: Cardiovascular Genetics* **7**, 771–781.

Tan BH, Valdivia CR, Rok BA, Ye B, Ruwaldt KM, Tester DJ, Ackerman MJ & Makielski JC (2005). Common human SCN5A polymorphisms have altered electrophysiology when expressed in PQ1077 splice variants. *Heart Rhythm* **2**, 741–747.

Tan BH, Valdivia CR, Song C & Makielski JC (2006). Partial expression defect for the SCN5A missense mutation G1406R depends on splice variant background Q1077 and rescue by mexiletine. *American Journal of Physiology – Heart and Circulatory Physiology* **291**, H1822–H1828.

Tan HL, Bink-Boelkens MT, Bezzina CR, Viswanathan PC, Beaufort-Krol GC, van Tintelen PJ, van den Berg MP, Wilde AA & Balser JR (2001). A sodium-channel mutation causes isolated cardiac conduction disease. *Nature* **409**, 1043–1047.

Tan HL, Kupershmidt S, Zhang R, Stepanovic S, Roden DM, Wilde AA, Anderson ME & Balser JR (2002). A calcium sensor in the sodium channel modulates cardiac excitability. *Nature* **415**, 442–447.

Tarradas A, Selga E, Beltran-Alvarez P, Pérez-Serra A, Riuró H, Picó F, Iglesias A, Campuzano O, Castro-Urda V, Fernández-Lozano I et al. (2013). A novel missense mutation, I890T, in the pore region of cardiac sodium channel causes Brugada syndrome. *PLOS ONE* **8**, e53220.

Tester DJ, Valdivia C, Harris-Kerr C, Alders M, Salisbury BA, Wilde AA, Makielski JC & Ackerman MJ (2010). Epidemiologic, molecular, and functional evidence suggest A572D-SCN5A should not be considered an independent LQT3-susceptibility mutation. *Heart Rhythm* **7**, 912–919.

Tsurugi T, Nagatomo T, Abe H, Oginosawa Y, Takemasa H, Kohno R, Makita N, Makielski JC & Otsuji Y (2009). Differential modulation of late sodium current by protein kinase A in R1623Q mutant of LQT3. *Life Sciences* **84**, 380–387.

Valdivia CR, Tester DJ, Rok BA, Munger TM, Jahangir A, Makielski JC & Ackerman MJ (2004). A trafficking defective, Brugada syndrome-causing SCN5A mutation rescued by drugs. *Cardiovascular Research* **62**, 53–62.

Vatta M, Dumaine R, Antzelevitch C, Brugada R, Li H, Bowles NE, Nademanee K, Brugada J, Brugada P & Towbin JA (2002). Novel mutations in domain I of SCN5A cause Brugada syndrome. *Molecular Genetics and Metabolism* **75**, 317–324.

Viswanathan PC, Benson DW & Balser JR (2003). A common SCN5A polymorphism modulates the biophysical effects of an SCN5A mutation. *The Journal of Clinical Investigation* **111**, 341–346.

Wang C, Wang C, Hoch EG & Pitt GS (2011). Identification of novel interaction sites that determine specificity between fibroblast growth factor homologous factors and voltage-gated sodium channels. *Journal of Biological Chemistry* **286**, 24253–24263.

Wang DW, Crotti L, Shimizu W, Pedrazzini M, Cantu F, De Filippo P, Kishiki K, Miyazaki A, Ikeda T, Schwartz PJ et al. (2008). Malignant perinatal variant of long-QT syndrome caused by a profoundly dysfunctional cardiac sodium channel. *Circulation: Arrhythmia and Electrophysiology* **1**, 370–378.

Wang DW, Desai RR, Crotti L, Arnestad M, Insolia R, Pedrazzini M, Ferrandi C, Vege A, Rognum T, Schwartz PJ et al. (2007). Cardiac sodium channel dysfunction in sudden infant death syndrome. *Circulation* **115**, 368–376.

Wang DW, Viswanathan PC, Balser JR, George AL & Benson DW (2002). Clinical, genetic, and biophysical characterization of SCN5A mutations associated with atrioventricular conduction block. *Circulation* **105**, 341–346.

Wang DW, Yazawa K, George AL & Bennett PB (1996). Characterization of human cardiac Na+ channel mutations in the congenital long QT syndrome. *Proceedings of the National Academy of Sciences* **93**, 13200–13205.

Wang HG, Zhu W, Kanter RJ, Silva JR, Honeywell C, Gow RM & Pitt GS (2016). A novel NaV1.5 voltage sensor mutation associated with severe atrial and ventricular arrhythmias. *Journal of Molecular and Cellular Cardiology* **92**, 52–62.

Wang L, Meng X, Yuchi Z, Zhao Z, Xu D, Fedida D, Wang Z & Huang C (2015). De novo mutation in the SCN5A gene associated with Brugada syndrome. *Cellular Physiology and Biochemistry* **36**, 2250–2262.

Wang SY, Tikhonov DB, Mitchell J, Zhorov B & Wang GK (2007). Irreversible block of cardiac mutant Na+ channels by batrachotoxin. *Channels* **1**, 179–188.

Watanabe H, Yang T, Stroud DM, Lowe JS, Harris L, Atack TC, Wang DW, Hipkens SB, Leake B, Hall L et al. (2011). Striking in vivo phenotype of a disease-associated human SCN5A mutation producing minimal changes in vitro. *Circulation* **124**, 1001–1011.

Wedekind H, Smits JP, Schulze-Bahr E, Arnold R, Veldkamp MW, Bajanowski T, Borggrefe M, Brinkmann B, Warnecke I, Funke H et al. (2001). De novo mutation in the SCN5A gene associated with early onset of sudden infant death. *Circulation* **104**, 1158–1164.

Wehrens XH, Rossenbacker T, Jongbloed RJ, Gewillig M, Heidbüchel H, Doevendans PA, Vos MA, Wellens HJ & Kass RS (2003). A novel mutation L619F in the cardiac Na+ channel SCN5A associated with long-QT syndrome (LQT3): a role for the I-II linker in inactivation gating. *Human Mutation* **21**, 552–552.

Winkel BG, Larsen MK, Berge KE, Leren TP, Nissen PH, Olesen MS, Hollegaard MV, Jespersen T, Yuan L, Nielsen N et al. (2012). The prevalence of mutations in KCNQ1, KCNH2, and SCN5A in an unselected national cohort of young sudden unexplained death cases. *Journal of Cardiovascular Electrophysiology* **23**, 1092–1098.

Yang P, Kanki H, Drolet B, Yang T, Wei J, Viswanathan PC, Hohnloser SH, Shimizu W, Schwartz PJ, Stanton M et al. (2002). Allelic variants in long-QT disease genes in patients with drug-associated torsades de pointes. *Circulation* **105**, 1943–1948.

Ye B, Valdivia CR, Ackerman MJ & Makielski JC (2003). A common human SCN5A polymorphism modifies expression of an arrhythmia causing mutation. *Physiological Genomics* **12**, 187–193.

Yokoi H, Makita N, Sasaki K, Takagi Y, Okumura Y, Nishino T, Makiyama T, Kitabatake A, Horie M, Watanabe I et al. (2005). Double SCN5A mutation underlying asymptomatic Brugada syndrome. *Heart Rhythm* **2**, 285–292.

Young KA & Caldwell JH (2005). Modulation of skeletal and cardiac voltage-gated sodium channels by calmodulin. *The Journal of Physiology* **565**, 349–370.

Zeng Z, Zhou J, Hou Y, Liang X, Zhang Z, Xu X, Xie Q, Li W & Huang Z (2013). Electrophysiological characteristics of a SCN5A voltage sensors mutation R1629Q associated with Brugada syndrome. *PLOS ONE* **8**, e78382.

Zhang J, Chen Y, Yang J, Xu B, Wen Y, Xiang G, Wei G, Zhu C, Xing Y & Li Y (2015). Electrophysiological and trafficking defects of the SCN5A T353I mutation in Brugada syndrome are rescued by alpha-allocryptopine. *European Journal of Pharmacology* **746**, 333–343.
